# Advanced physiological maturation of iPSC-derived human cardiomyocytes using an algorithm-directed optimization of defined media components

**DOI:** 10.1101/2022.10.10.507929

**Authors:** Neal I. Callaghan, Lauren J. Durland, Wenliang Chen, Uros Kuzmanov, Maria Zena Miranda, Zahra Mirzaei, Ronald G. Ireland, Erika Yan Wang, Karl Wagner, Michelle M. Kim, Julie Audet, J. Paul Santerre, Anthony O. Gramolini, Filio Billia, Milica Radisic, Seema Mital, James Ellis, Peter H. Backx, Craig A. Simmons

## Abstract

Induced pluripotent stem cell-derived cardiomyocytes (iPSC-CMs) hold tremendous promise for in vitro modeling to assess native myocardial function and disease mechanisms as well as testing drug safety and efficacy. However, current iPSC-CMs are functionally immature, resembling in vivo CMs of fetal or neonatal developmental states. The use of targeted culture media and organoid formats have been identified as potential high-yield contributors to improve CM maturation. This study presents a novel iPSC-CM maturation medium formulation, designed using a differential evolutionary approach targeting metabolic functionality for iterative optimization. Relative to gold-standard reference formulations, our medium significantly matured morphology, Ca^2+^ handling, electrophysiology, and metabolism, which was further validated by multiomic screening, for cells in either pure or co-cultured microtissue formats. Together, these findings not only provide a reliable workflow for highly functional iPSC-CMs for downstream use, but also demonstrate the power of high-dimensional optimization processes in evoking advanced biological function in vitro.

## Introduction

Cardiovascular diseases (CVDs) including heart failure, myocardial infarction, and arrhythmias often have multifactorial etiologies, are progressive and prone to sequelae, and cannot usually be curatively treated; for this reason, they form the single most common cause of death in developed countries ^1^. For decades, knowledge on CVDs has been advanced using animal-derived models of function and disease, but the need for robust and accurate *in vitro* models to inform human-specific pathophysiology has become apparent ^2^. Similarly, the incidence of cardiotoxicity, which has remained unpredicted by preclinical *in vivo* screening with murine models, is a leading reason for failure of new drug trials or post-marketing drug recalls ^3,4^. iPSC-CMs provide a promising avenue for cardiac pharmaceutical efficacy screening, both for preclinical screening and personalized efficacy testing ^5–7^. Both disease phenotypes and drug responses are emergent composites of interactions across multiple aspects of physiology ^8,9^, necessitating higher-fidelity *in vitro* models of myocardium. However, current iPSC-CMs are phenotypically immature in all aspects of CM-specific physiology; they do not recapitulate the contractile force, morphology, electrophysiology, calcium handling characteristics, or metabolic profile associated with adult myocardium ^10^, and are not satisfactorily predictive of patient responses to pharmaceuticals ^11^.

Culture medium optimization has been explored as a high-yield avenue for further iPSC-CM maturation. Notable advances have been made using one- or two-factor experiments including metabolic substrates such as fatty acids or galactose ^12–14^, and hormones such as triiodothyronine (T3) ^15–20^, insulin ^21^ or insulin-like growth factor ^16,22^, glucocorticoids ^16,20,21^, or the B27 supplement comprising a mixture of hormones and small molecules optimized for functional neuronal culture ^13,15,17,23–26^. However, there has been comparatively little focus to date on supplementation of endogenous cofactors or otherwise functional small molecules. The energetic demands of a fully mature CM to fuel electrophysiological and contractile function (e.g., via oxidative phosphorylation) may preclude secondary metabolic pathways that would produce such molecules from more basal media. Furthermore, pairwise and higher-order interactions have been found to be ubiquitous across biological systems such as cell fate, comprising differentiation or maturation and their associated homeostatic changes ^27,28^. As a result, small stepwise optimization experiments are unlikely to converge, even in aggregate, on high-efficacy maturation medium formulations.

In this study, we sought to mature iPSC-CMs using a suite of soluble factors comprising metabolic substrates, hormones, cofactors, and other small molecules based on the potential for widescale interactions between factors. Our general strategy in choosing factors was to mirror the exogenous availability of soluble signals to which the myocardium is exposed during late cardiac development, specifically during the perinatal window and early childhood ^29^, developmental phases that coincide with a switch to a predominantly oxidative metabolism ^30^. This metabolic specialization prioritizes oxidative catabolic flux and outsources peripheral metabolic processes (e.g,. anaerobic catabolism, glycogen-related processes, cofactor synthesis, etc.) to other tissues. This transition to a highly efficient energetic phenotype may drive maturation in other functional metrics by increasing the accessible short-term energetic capacity within the cell ^31^. To maximize factor synergies and high-order interactions, a high dimensional, differential evolution (HD-DE) algorithm was employed to functionally interrogate a large factor space, even in the presence of factor interactions or nonlinear responses ^32^. By combining a set of 17 independent soluble factors each at 5 dosage levels, a traditional full factorial screen would require c.a. 763 billion runs to cover the solution space; using the HD-DE methodology, we developed a high-performing custom formulation after 4 iterative generations comprising 169 unique formulations. Candidate formulations were scored based on their oxidative uncoupled:control ratio using cell respirometry as a self-normalizing correlate of overall functional maturation. This approach identified a medium formulation that best enhances iPSC-CM maturity to maximize multiple functional performance metrics compared to existing gold-standard commercial and published formulations. Our formulation demonstrated an unmatched degree of advancement of many specific functional metrics of maturity including morphology, contractility, electrophysiology, Ca^2+^ handling, metabolism, and gene and protein expression profiles.

## Results

### Differential evolution and selection of an iPSC-CM maturation medium

To efficiently optimize iPSC-CM maturation medium without full mechanistic understanding of the processes involved, we implemented an HD-DE algorithm-driven process (**Figure 1a**), using oxidative uncoupled:control ratio (UCR; the ratio of pharmaceutically-uncoupled to steadystate oxygen consumption rates) as an objective metric to score successive generations of candidate formulations relative to a gold-standard control (STEMCELL Cardiomyocyte Maintenance Medium) (**Figure 1b**). A total of 17 soluble factors (**Table S1, Figure 1c**) were chosen for iterative optimization; 6 of these factors as well as M199 basal medium had not previously been tested, nor appreciable analogues, for their use in CM maturation. Factors were surveyed at any of 5 discrete doses (including null) within a given candidate formulation. In all, 169 unique formulations were tested over 204 runs across 4 iterative generations each consisting of 3 weeks of treatment with candidate formulations (culture timelines in **Figure S1a-b**). By principal component analysis (PCA), the top 10 performing formulations formed 2 main clusters of composition, with 2 outliers (**Figure 1d-e**); substrates and cofactors were correlated most strongly in PC1, while hormones and small molecules primarily associated with signaling tended to align with PC2. A final formulation for further characterization was manually defined by including specific dose-factor combinations present in the top-right cluster of PCA but absent from the bottom-left quadrant, resulting in a composite of high-performing candidate formulations. This 16-component iPSC-CM maturation formulation (“C16”) was confirmed to be a viable and high-performing candidate against gold-standard formulations from STEMCELL, iCell, and the homemade RPMI + B27 cocktail, using both morphological and contractile comparisons after 3 weeks of culture. iPSC-CM monolayers treated with C16 medium demonstrated marked cell hypertrophy, anisotropy, and spontaneous alignment (**Figure S1c**), as well as clear sarcomeric striations under phase-contrast in C16-treated monolayers (**Figure S1d-e**). Confocal microscopy demonstrated that C16 medium enhanced myofibrillar density, α-actinin striation, and Cx43 junctional expression (**Figure 1f**), and induced striated expression of hallmark CM functional proteins DHPR, NCX (**Figure 1g**), SERCA2a and RyR2 (**Figure 1h**). Optical contractile traces, obtained using particle image velocimetry, demonstrated accelerated full contractile cycles in C16-cultured monolayers derived from two iPSC-CM lines from healthy donors ^33^ (PGPC17 and PGPC14), as well as pathologically slowed and biphasic contraction in a *MYBPC3-*KO CRISPR-generated PGPC17 mutant iPSC line (**Figure S1f**).

**Figure 1.**
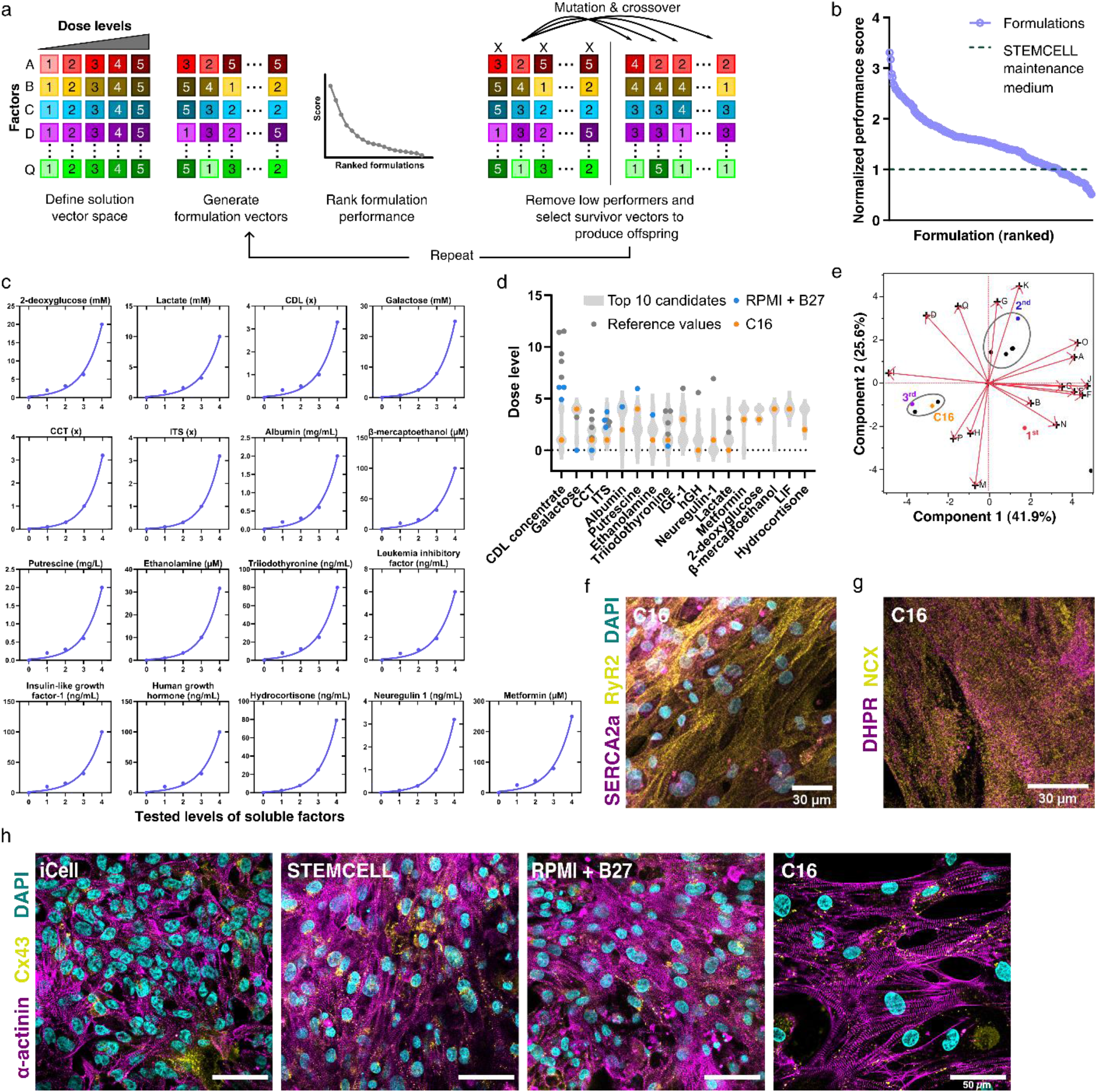
Iterative development and performance of iPSC-CM maturation medium candidates relative to gold-standard formulations. **a)** Visual representation of High Dimensional, Directed Evolution (HD-DE) algorithm workflow; from a quasi-random distribution of vectors (formulations) in an extensive but discrete solution space, low-performing candidates are discontinued while high-performing candidates are kept and iterated using mutation and crossover operations. **b)** Ranked performance of all tested formulation candidates (n =169 unique vectors over 204 instances) against STEMCELL Technologies Cardiomyocyte Maintenance Medium (dashed line), using respirometric uncoupled-control ratio as an objective metric. **c)** Soluble additive factors and their tested levels included in the HD-DE optimization. **d)** Final composition of C16 iPSC-CM maturation medium by factor (orange), superimposed over factor levels of the top 10 formulation candidates in the optimization phase (grey violin plots), and analogous reference levels, where applicable, from the literature (dark grey) or in the commonly used iPSC-CM culture medium RPMI + B27 (blue). **e)** Principal component analysis biplot visualization of the top ten global performing formulations during optimization (points; red indicates 1^st^ place formulation, blue indicates 2^nd^ place formulation, and purple indicates 3^rd^ place formulation) in addition to a composite, manually-defined formulation (“C16”, gold); red vector arrows represent correlation of medium additives (A-Q) corresponding to the order of the additives in **Table S1**. Circled formulation groups indicate high-performing clusters derived from common lineages over generations of HD-DE optimization. **f)** Confocal detail (DAPI-stained nuclei represented in cyan) of sarco/endoplasmic reticulum Ca2+ ATPase 2a (SERCA2a, magenta) and cardiac ryanodine receptor (RyR2, yellow) localization in C16-treated monolayer. **g)** Confocal detail of DHPR (magenta) and sodium-calcium exchanger (NCX, yellow) localization in C16-treated monolayer. All analyses performed after 3 weeks of medium treatment. **h)** Confocal images of α-actinin(magenta) and Cx43 (yellow)-stained monolayers (DAPI-stained nuclei represented in cyan) cultured in C16, iCell, RPMI + B27, or STEMCELL media; scale bars 50 μm. All analyses performed after 3 weeks of medium treatment.

### Electrophysiological characterization and I_K1_ identification in maturing iPSC-CMs

The presence of stable resting (diastolic) membrane potentials and the absence of spontaneous beating are hallmarks of mature working myocardium. Importantly, in addition to the maturation properties already listed, iPSC-CM monolayers treated with C16 medium were not spontaneously contractile, unlike RPMI + B27-treated cells, suggesting marked electrophysiological differences between treatments. To assess the electrical properties of our cultures, we measured intracellular voltages in cells after 6 weeks of treatment with either C16 or RPMI + B27 (**Figure 2a**). These electrical differences were associated with classical differences in the resting diastolic membrane potentials. Specifically, RPMI + B27 cells underwent spontaneous depolarization after reaching minimum diastolic membrane potentials (MDPs) of −56.1 ± 6.8 mV (**Figure 2b**), a pattern that is characteristic of immature cardiomyocytes, while C16-treated cells displayed fixed resting diastolic membrane potentials (−81.1 ± 7.7 mV). RPMI + B27-treated cell APs were accompanied simultaneously by contraction and Ca^2+^ transients (**Figure 2c**) at a rate of ∼ 0.2 Hz. By contrast, C16-treated cells did not display spontaneous action potentials (APs), beating or Ca^2+^ transients in the absence of electrical stimulation. Previous studies have established that the absence of spontaneous APs in mature tissues is linked to the presence of background inward rectifier K^+^ channels (I_k1_), which are able to “clamp” the resting membrane potential at values close to the equilibrium potential for K^+^ ions (E_K_) ^34^. To assess whether I_K1_ underlies the differences in spontaneous APs and beating between C16 and RPMI + B27, we applied extracellular Ba^2+^ at a dose (400 μmol L^−1^) that potentially blocks I_K1_ channels. While Ba^2+^ addition had minimal effect on either MDPs (49.2 ± 5.3 mV) or the beating rates of RPMI + B27 cells, it caused C16-treated cells to display spontaneous beating and APs accompanied by spontaneous depolarizations from MDPs of −53.7 ± 1.8 mV. Our findings suggest that C16treated cells express high number of I_K1_ channels compared to RPMI + B27-treated cells. Indeed, in voltage-clamp studies, C16-treated cells demonstrated robust background K^+^ currents with characteristic kinetic behaviour, classical inward rectification properties (**Figure 2d-e**) and reversal properties (i.e., ∼E_K_) which are hallmark features of I_K1_. Moreover, these background currents in the C16 treatment were eliminated by 400 μmol L^−1^ Ba^2+^ (**Figure 2e**). RPMI + B27 cells also displayed Ba^2+^-sensitive background K^+^ currents but these were ∼4-fold smaller than in C16-treated cells (*i*.*e*., Ba^2+^sensitive densities measured at −115 mV were −21.9 ± 8.2 pA pF^−1^ in C16 versus −5.5 ± 3.9 pA pF^−1^ in the RPMI + B27 treatment, **Figure 2f**), consistent with the RPMI + B27 treatment, **Figure 2f**), consistent with the observed effects of Ba^2+^ on APs and beating of C16-*vs*. RPMI + B27-treated cells.

**Figure 2.**
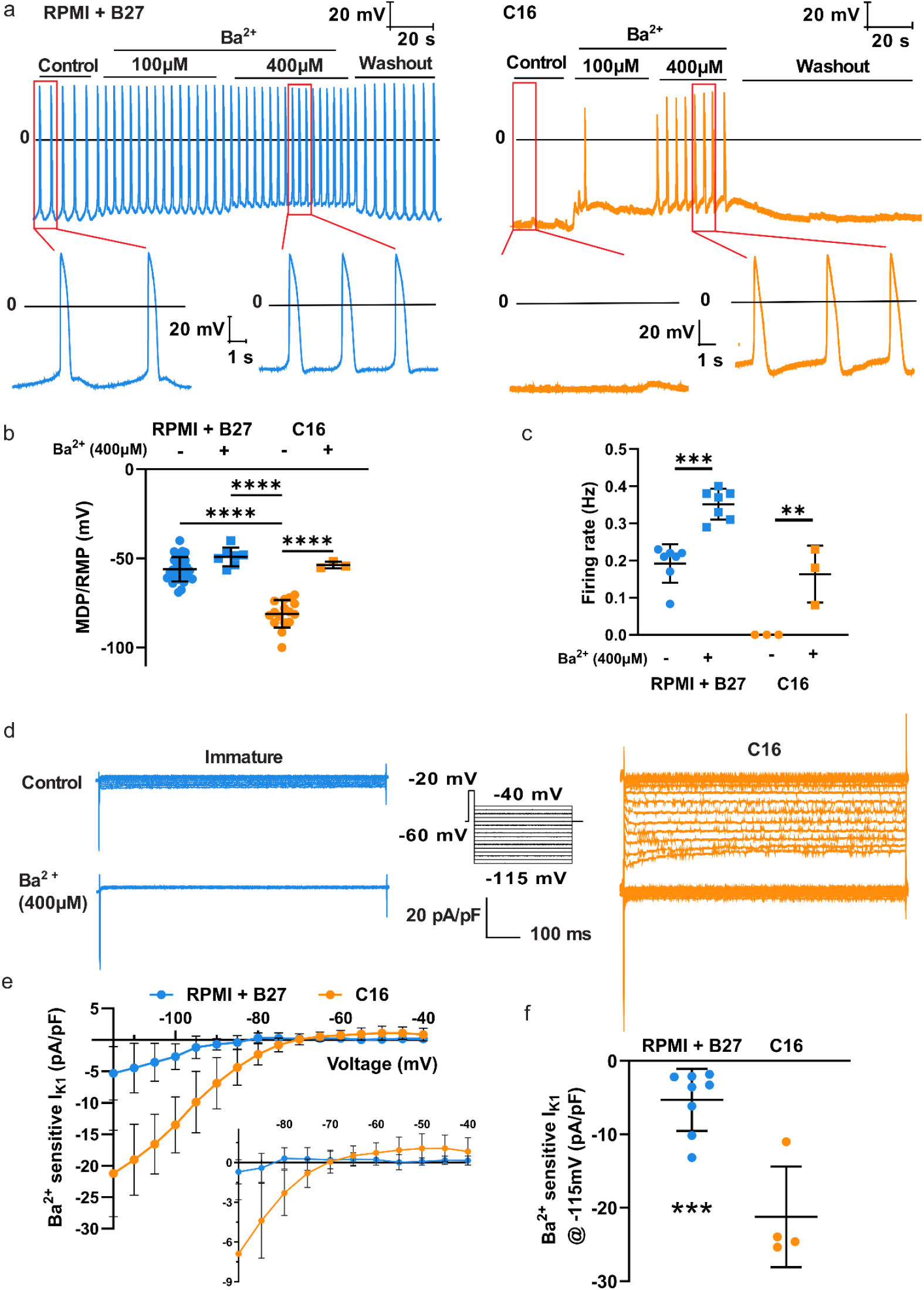
Electrophysiological enhancement of iPSC-CMs matured using C16 medium. **a)** Sample traces of intracellular electrode recordings of iPSC-CMs treated with C16 maturation medium (left) or RPMI + B27 (right) for 6 weeks, in neat Tyrode’s solution, followed by sequential supplementation with 100 and 400 μmol L^−1^ BaCl_2_, before eventual washout. **b)** Minimum diastolic potentials (resting membrane potentials) from intracellular recording in cell sheets, with or without 400 μmol L^−1^ BaCl_2_ perfusion. **c)** Spontaneous action potential firing rates measured during action potential recordings, with or without 400 μmol L^−1^ BaCl_2_ perfusion. **d)** Inward rectifier (I_K1_) currents in neat Tyrode’s and supplemented with 400 μmol L^−1^ BaCl_2_; inset indicates voltage protocol and scale. **e)** Voltage sensitivity curves of IK1; inset demonstrates positive component of I_K1_ at high voltage. **f)** I_K1_ current density at −115 mV. Data points denote biological replicates. Error bars denote mean ± SD; *, **, ***, and *** denote p < 0.05, 0.01, 0.001 and 0.0001, respectively.

### Physiological characterization of maturing iPSC-CMs

iPSC-CM monolayers treated with C16 medium demonstrated differential Ca^2+^ handling and metabolic flux compared to gold-standard formulations after 6 weeks of culture. Measurement of steady-state Ca^2+^ handling at 1 Hz pacing using line-scanning microscopy demonstrated accelerated onset (**Figure 3a-b**) and decay kinetics (**Figure 3c-e**) in C16-treated cells vs. controls, which may reflect a greater dependence of Ca^2+^ cycling on the sarcoplasmic reticulum (SR) seen in mature versus immature myocardium ^10^. Consistent with increased SR contributions, administration of the β-adrenergic receptor agonist, isoproterenol, further accelerated Ca^2+^ transient onset in C16-treated cells only (**Figure 3f**). More direct evidence of enhanced SR function in the C16 culture conditions was uncovered by perfusing cells with high levels of caffeine to cause complete release of the SR’s Ca^2+^ into the cytosol. Specifically, when cells were pre-treated with verapamil to prevent Ca^2+^ entry into the cytosol via L-type channels, a bolus of caffeine generated a rise in intracellular Ca^2+^ within C16 cells that was ∼2-fold higher than for cells cultured in STEMCELL medium (p < 0.0001) (**Figure 3g-i**). C16-treated iPSC-CMs also diverged from gold-standard formulations in metabolic profiling using Seahorse XFe96 respirometry (**Figure 3j-k**). Specifically, the per-cell basal oxygen consumption rate outpaced control treatments (**Figure 3l**), while the FCCP-enabled UCR of C16-cultured cells was higher than those of the iCell (p = 0.035), RPMI + B27 (p = 0.023), and STEMCELL (p = 0.013) control treatments (**Figure 3m**). These findings were consistent with targeted metabolomic analyses of iPSC-CM monolayers at the same timepoint, treated with either the C16 formulation, a leading high-efficacy formulation by Feyen *et al*. ^35^, or RPMI + B27 as a control. C16-treated monolayers demonstrated greater incorporation (*i*.*e*., lowered m+0 proportion) of labeled glucose and lactate into aerobic pathways (**Figure 3n-q**), with reduced anaerobic metabolism (**Figure 3r-t**) while maintaining pentose phosphate flux (**Figure 3u**).

**Figure 3.**
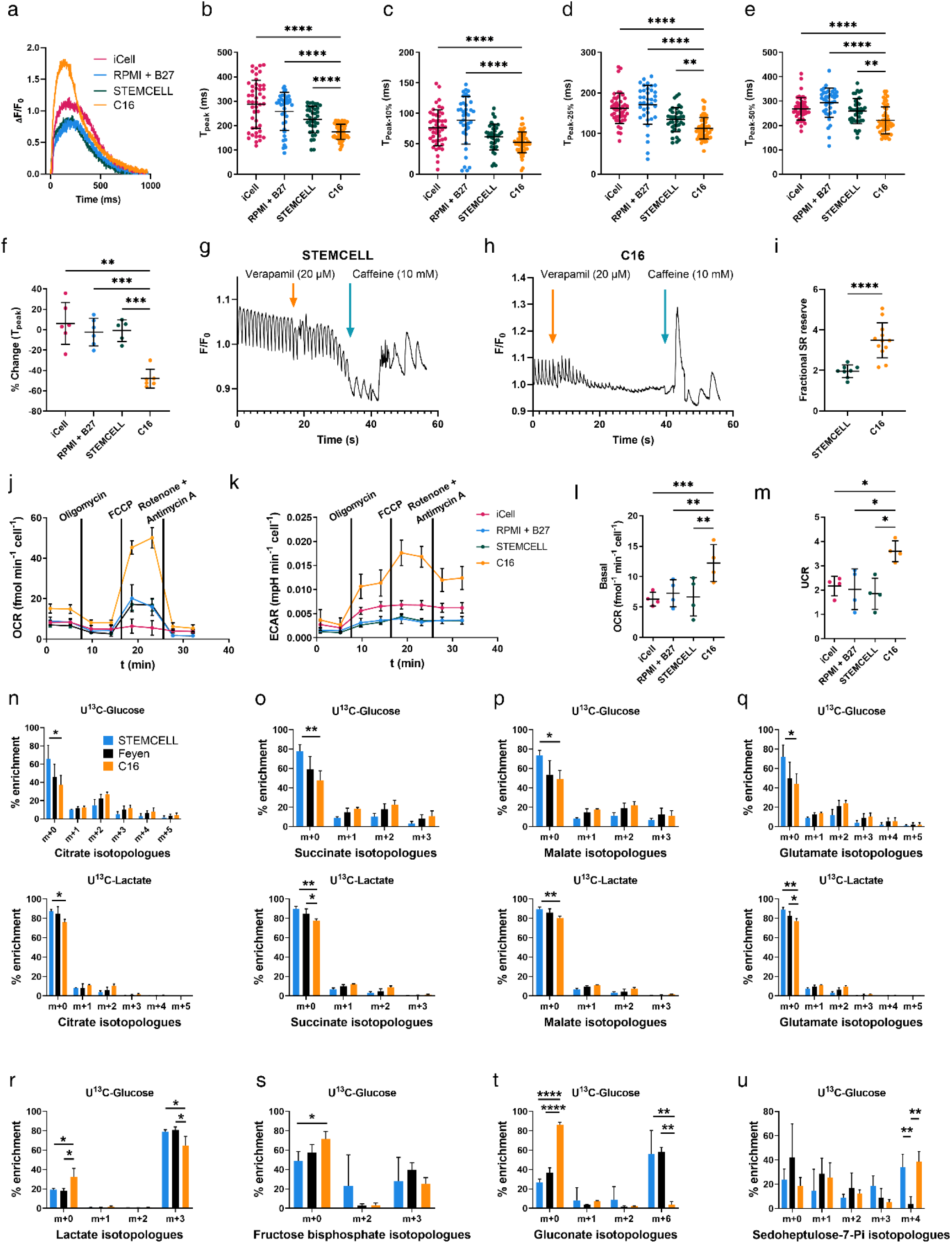
Functional enhancement of iPSC-CMs matured in C16 medium. **a)** Representative line-scanning Fluo-4 traces during steady-state Ca^2+^ transients at 1 Hz pacing of single iPSC-CMs treated 6 weeks with C16 maturation medium compared to gold-standard iPSC-CM media (iCell Cardiomyocytes Maintenance Medium, RPMI + B27, and STEMCELL Cardiomyocyte Maintenance Medium); kinetic metrics isolated from trace sets include time to peak **(b)**, and times to 10% **(c)**, 25% **(d)**, and 50% **(e)** decay from peak fluorescence, respectively.. **f)** Relative change (%) of baseline time to peak fluorescence in single cells upon addition of isoproterenol (5 μmol L^−1^). Fluo-4enabled Ca^2+^ traces of C16 **(g)** and STEMCELL **(h)**-treated cultures after 6 weeks with steady-state pacing at 1 Hz followed by sequential additions of verapamil (20 μmol L^−1^) and caffeine (10 mmol L^−1^). **i)** Caffeine-induced Ca^2+^ unloading amplitudes normalized to steady state transient amplitudes. Representative mitochondrial stress test traces demonstrating **(j)** oxygen consumption (OCR) and **(k)** extracellular acidification (ECAR) rates of iPSCCMs cultured 6 weeks in C16 medium or existing gold-standard control formulations, normalized to cell number. Normalized basal oxygen consumption rates **(l)** and uncoupled-control ratios **(m)** by treatment. Enrichment of central carbon metabolic pathways **(n-q)** and peripherally-associated metabolite pools **(r-u)** with either uniformly ^13^C-labeled glucose or lactate; including citrate **(n)**, succinate **(o)**, malate **(p)**, glutamate **(q)**, lactate **(r)**, fructose-1,6-bisphosphate **(s)**, sedoheptulose-7-phosphate **(t)**, and gluconate **(u)**. Error bars denote mean ± SD; *, **, *** and **** denote p < 0.05, 0.01, 0.001 and 0.0001, respectively.

### Transcriptomic profiling of maturing iPSC-CMs

RNA sequencing was used to evaluate the transcriptomic profiles of iPSC-CMs treated with C16 or gold standard maintenance media to help identify targets for further physiological analysis. PCA was used to compare the overall transcriptional profiles of each medium relative to immature day 20 (d20) controls, demonstrating that C16-treated cells were highly divergent from both controls and other media (**Figure 4a**). Many of the GO terms significantly upregulated in C16 treatments were associated with general cardiomyocyte functions, including the cardiac action potential, contraction, regulation of contractility, and oxidative metabolism, while GO terms heavily downregulated in C16 treatments included differentiation, motility, and anaerobic metabolism. Similarly, gene set enrichment analysis (GSEA) comparing C16 to gold-standard control treatments revealed GO terms pertaining to heart function, lipid metabolism and oxidative phosphorylation, cell-cell adhesion, and transcriptional regulation (**Figures S2-3**).

**Figure 4.**
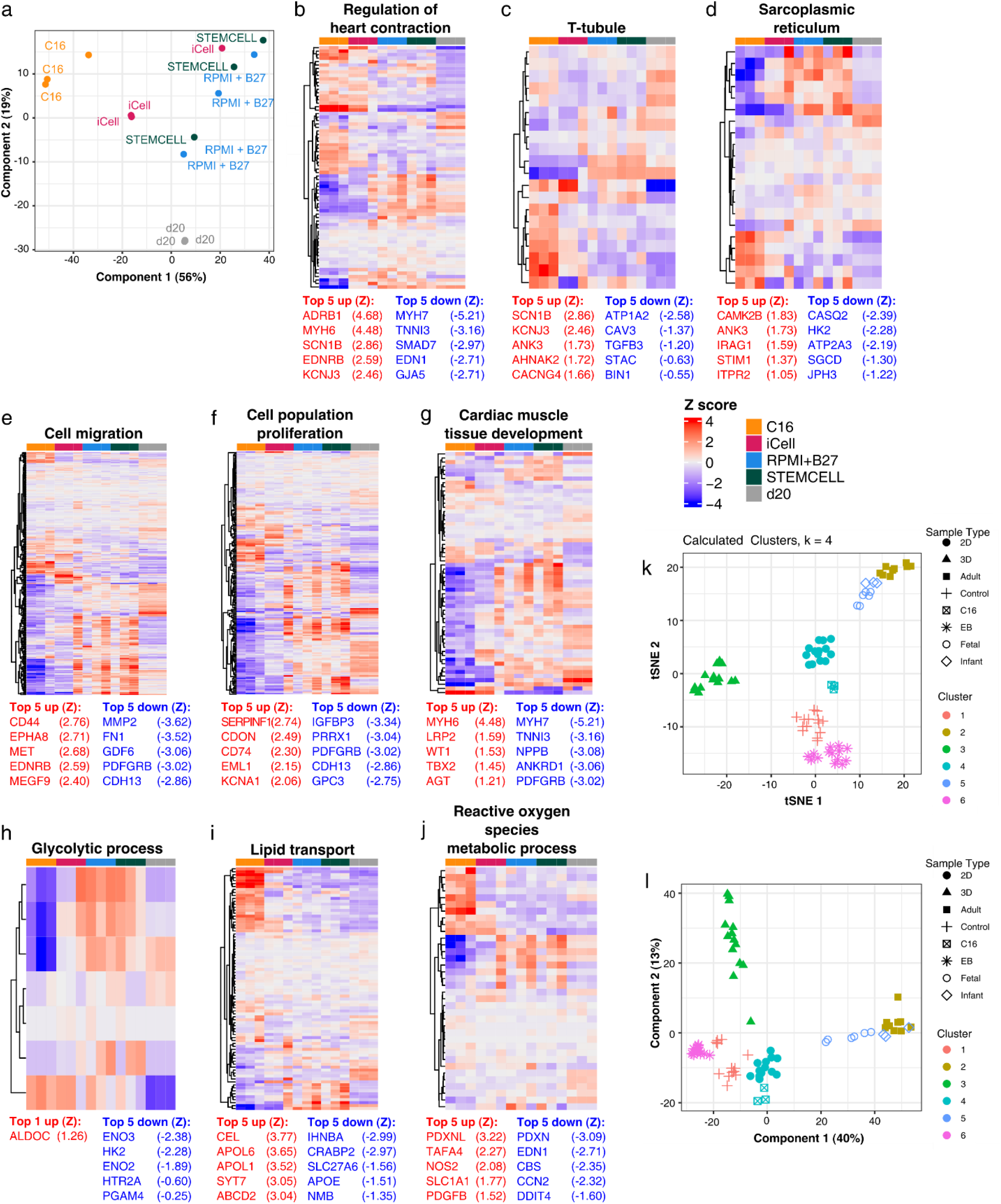
Transcriptomic profiling of matured iPSC-CMs and RNA sequencing-enabled meta-analysis to ex vivo tissues and historical maturation expriments. **a)** Principal Component analysis of iPSC-CoMs cultured in C16 medium or a gold-standard formulation for 6 weeks, or at d20 of differentiation and prior to treatment (control). Colours denote different sample groups. **b-j)** Heatmaps displaying expression of select genes from the GO terms; top 5 upregulated (red) and downregulated (blue) genes within C16-treated cells per GO term listed with their respective Z scores. n = 3 biological replicates per treatment. Head-to-head treatment comparisons found in **Figures S2-S3. k-l)** Meta-analysis of bulk RNA sequencing of iPSC-CM in vitro maturation compared to native CMs at fetal, infant, and adult life stages, including tSNE **(k)** and principal component analysis **(l)**. Sample descriptions found in **Table S2**; validation in **Figure S4**, principal component attribution in **Figure S5**, and cluster analysis in **Figure S6**.

### Meta-analysis of (iPSC-)CM maturation

RNA-seq datasets from several studies of CM maturation were pooled into a single meta-analysis to better understand transcriptional trajectories, or the gene expressional landscape around the physiological maturation process. To enable interpretation, external samples were divided into one of 7 broad categories. *In vivo* myocardial or CM samples were assigned to the ‘Adult’, ‘Infant’ or ‘Fetal’ category based on developmental timepoint at isolation. Matured *in vitro* iPSC-CM samples were assigned to the “2D”, “3D” or “EB” categories, depending on the culture methodology and type of maturation approach employed (see **Table S2** for detailed descriptions). ‘Control’ samples were immature iPSC-CMs, collected at ∼d18-20 of differentiation, after the onset of spontaneous beating but prior to starting any type of maturation protocol.

To adjust for batch effects introduced by the inclusion of multiple datasets, the count matrix was adjusted to minimize variation among samples from different studies, while preserving variation among the designated sample categories (**Figure S4**). Following batch correction, tSNE and PCA analyses demonstrated that the external samples could agnostically be re-separated into distinct clusters according to their pre-assigned categories (**Figure 4k-l**). Both GSEA of principal components (**Figure S5**) and differential expression analysis (**Figure S6**) revealed similar trends in functional gene expression between unique clusters; the PC1 positive direction (2D cluster) was enriched for electrophysiological function and electrical conduction, PC2 positive (3D cluster) enriched for contractile function and Ca^2+^ handling, and PC1 negative (EB cluster) enriched for cardiac developmental and morphogenic processes. DEA further identified specific functions enriched in each cluster that were not identified according to the PCA loadings, such as GPCR signaling and lipid handling in the “3D” cluster, and Ca^2+^ handling in the “2D” cluster.

### Proteomic profiling of maturing iPSC-CMs

Transcriptomic analysis suggested marked changes to transcriptional regulation, protein synthesis and turnover, and protein trafficking. To directly examine protein changes in the 2D system, global proteomic analysis of urea-soluble fractions was conducted on iPSC-CMs after 6 weeks treatment in C16, iCell, RPMI + B27, and STEMCELL media, as well as the formulation designed by Feyen *et al*. ^35^ (**Figure 5**). Principal component analysis (PCA), conducted on a set 1869 proteins that had valid (greater than 0 LFQ intensity) values in all three samples from at least one experimental treatment, demonstrated that PC1 largely separated the STEMCELL and C16 conditions, while PC2 separated the high-performing Feyen medium from the other four conditions (**Figure 5a**). Both PC loadings included metabolic and mitochondrial-specific proteins; the positive direction of PC1 also heavily featured sarcoplasmic synthesis and maintenance ontologies, ionoregulatory proteins, and membrane trafficking components, while the positive direction of PC2 heavily featured cytoskeletal proteins associated with contractility. Enriched GO terms associated with maturation conditions included membrane functionalization, oxidative metabolism, and mRNA processing, while downregulated terms centred on apoptotic induction and protein localization (**Figure 5b-c**). Of particular note, MYH7 protein expression was not significantly downregulated in C16-treated samples, in contrast with its transcriptional downregulation in identical samples. This may suggest that regulation beyond the level of mRNA transcription may contribute to the functional expression of maturity-associated protein products.

**Figure 5.**
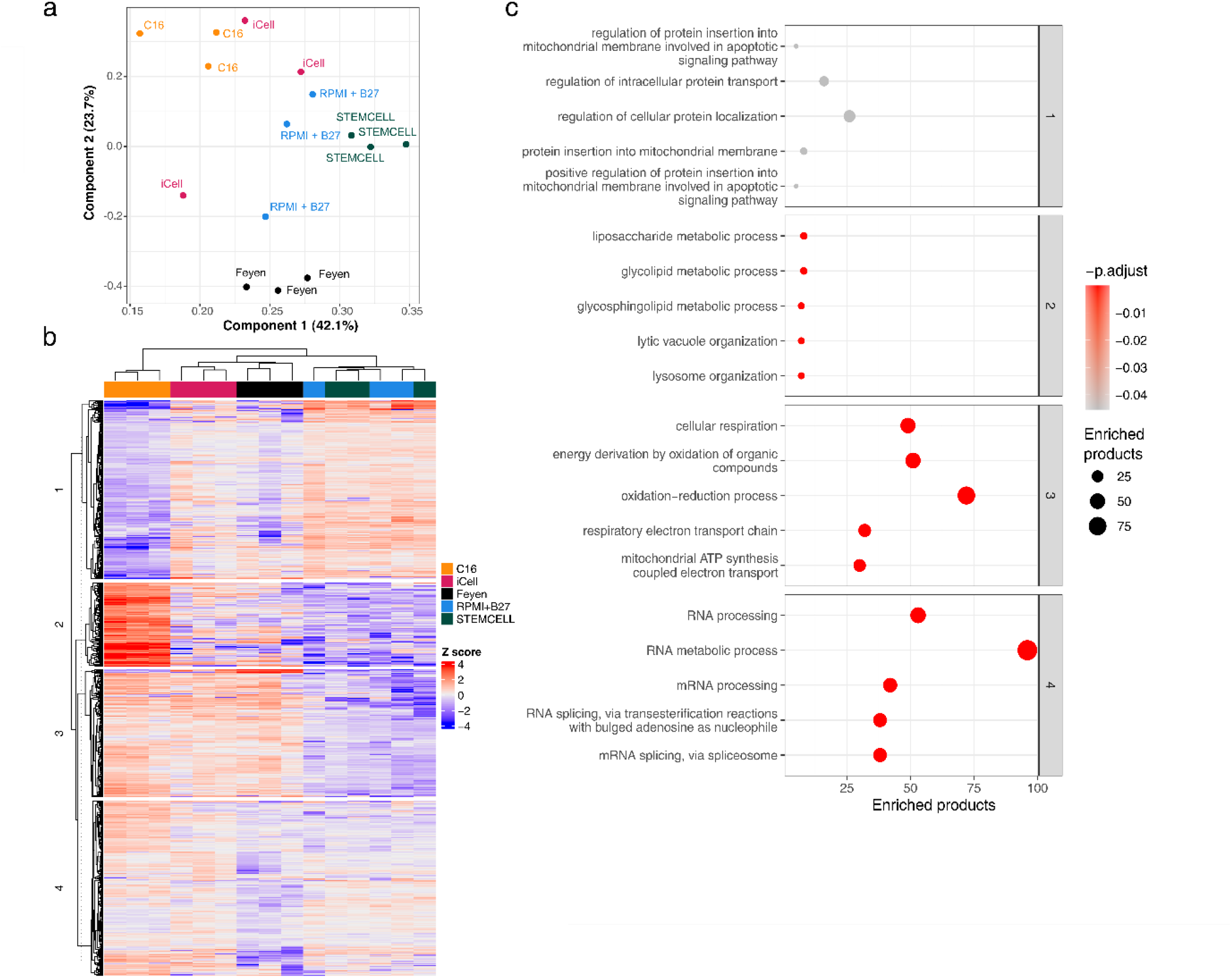
Global proteomic profiling of C16-treated iPSC-CMs relative to gold-standard formulations. **a)** Principal component analysis of differential protein expression in iPSC-CMs after 6 weeks of treatment in C16, iCell, RPMI + B27, or STEMCELL media, as well as a top-performing formulation from the literature ^35^. **b)** Heat map demonstrating stratification of treatments by protein regulation. **c)** Over-representation analysis of the top 5 GO terms differentially expressed in C16-treated iPSC-CMs compared to other treatments for each cluster in panel **(b)**; GO term size (in number of gene products) represented both by marker size and size on X-axis.

### Maturation of iPSC-CM-containing engineered cardiac microtissues

Marked differences in the level of attainable iPSC-CM function relative to monolayers have previously been found in the context of engineered and co-cultured microtissues, leveraging physiological mechanical and electrical cues to drive maturation. We used the Biowire II platform ^36^ to examine further the impact of our maturation medium on parameters of CM function (**Figure 6**). These studies involved the creation of compacted multicellular muscle strips ^36^, followed by 3 weeks of culture in either modified C16 (see Methods) or RPMI + B27, and with or without continual electrical field pacing (1 Hz). As in monolayer culture, Biowire tissues in C16 media exhibited lower rates of spontaneous contractility than in RPMI + B27 (**Figure 6a**) after 3 weeks in of treatment. Contractile forces normalized to cross-sectional area (*i*.*e*., twitch/systolic stress) was ∼5fold higher with C16 versus RPMI + B27 treatment after 1, 2, or 3 weeks in culture (**Figure 6b**), with or without electrical stimulation during the culture period. After 3 weeks in culture with electrical stimulation, the twitch stress was 1.69 ± 1.42 mN mm^−2^ with C16 treatment *vs*. 0.35 ± 0.37 mN mm^−2^ with RPMI + B27 treatment. By 3-way ANOVA, C16 treatment (p = 0.0002) and culture time (p < 0.0001) both independently enhanced twitch stress, and these parameters positively interacted (p < 0.0001), establishing greater benefit of culture time in C16 medium versus the goldstandard control. Neither tissue morphology nor diastolic stresses were significantly affected by treatments (**Figure S7**). Furthermore, positive force-frequency responses after 3 weeks in culture were enhanced when measured at either 2 Hz or 3 Hz stimulation (relative to tissue-matched baseline stresses at 1 Hz) by both culture time (p < 0.0001) and C16 treatment (p < 0.0001) (**Figure 6c**); again, there was a positive interaction (p < 0.0001) between these factors.

**Figure 6.**
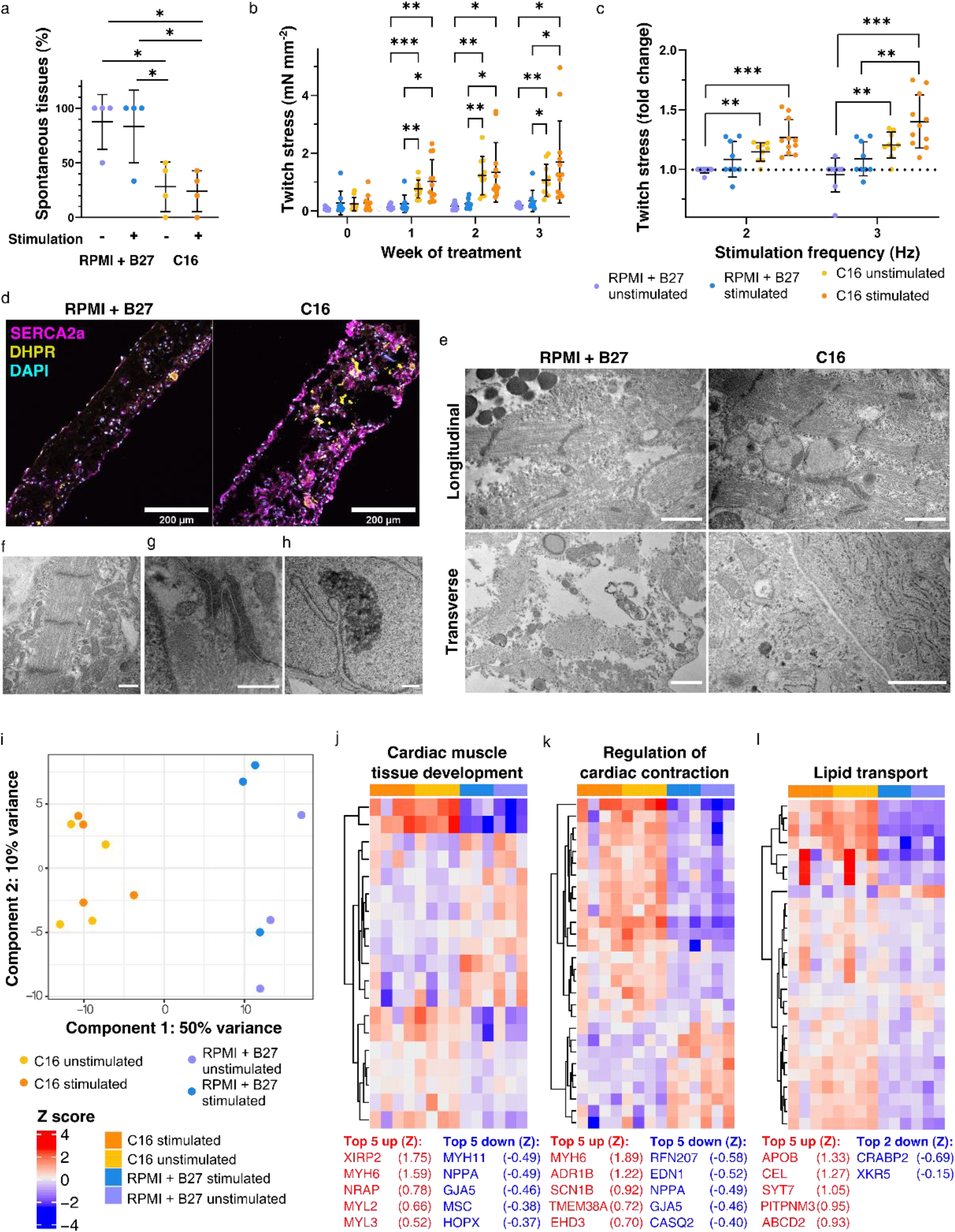
Contractile enhancement of engineered myocardial tissues cultured in C16 maturation medium. **a)** Decrease in tissue contractile spontaneity after 3 weeks of culture in C16 media. **b)** Progressive absolute twitch force increase over 3 weeks of Biowire II culture in C16 or RPMI + B27 medium formulations, with or without continuous electrical field stimulation (1 Hz). **c)** Positive force-frequency relationship of contractile displacement of polymeric beams summatively increases with both mediumand electrical stimulation-mediated maturation of tissues after 3 weeks of tissue treatment in C16 or RPMI + B27 media; measurements are normalized to 1 Hz stimulation of the relevant treatment condition (dotted line). **d)** Confocal microscopy reveals differential expression of SERCA2a (magenta) and DHPR (yellow) in electrically-stimulated Biowires… cultured 3 weeks in C16 or RPMI + B27 media. **e)** TEM of longitudinal and transverse sections of C16-treated Biowires induces sarcomeric development and higher ribosome-associated sarcoplasmic reticulum (scale bars 1 μm); detailed examination of C16-treated tissues (**e-h**; scale bars 500 nm) include sarcomeres with defined M- and Hbands and Z-discs and myofibril-associated high mitochondrial density (f), electrondense intercalated discs **(g)**, and nuclear SR invaginations **(h). i-l)** Transcriptomic analysis of Biowire II cultures treated with C16 or RPMI + B27 medium and with or without electrical field stimulation. Principal component analysis **(i)** and associated GO terms stratifying treatments top 5 upregulated (red) and downregulated (blue) genes within all C16-treated tissues per GO term listed with their respective Z scores; n = 4 biological replicates per C16 treatment and n = 3 biological replicates per RPMI + B27 treatment. Error bars denote mean ± SD. Significance indicated by * p < 0.05, ** p < 0.01, and *** p < 0.005. Stress value calibration and tissue characterization found in **Figure S7**; breakdown of principal components in **(i)** found in **Figure S8**.

Tissues treated for 3 weeks with the C16 formulation also exhibited higher-density SERCA2a and DHPR staining than those treated with RPMI + B27, similarly to in monolayers (**Figure 6d**). However, DHPR expression was variable between cells, suggesting stratification of phenotypes within the tissue. Compared to RPMI + B27 treatment, C16-treated tissues under transmission electron microscopy demonstrated increased myofibrillar bundling, sarcomeric striation and zonation, mitochondrial density and myofibrillar association, electron-dense intercalated discs, and nuclear invaginations by the SR ^37^ (**Figure 6e-h**). Consistent with both the functional and morphological differences observed in C16-treated tissues, the two formulations were highly stratified by PCA of bulk RNA sequencing primarily by medium treatment and not electrical field stimulation (**Figure 6i**). Highly enriched GO terms in C16-treated cultures pertained most strongly to oxidative metabolic and contractile ontologies, while RPMI + B27-treated culture segregation was driven by developmental ontologies (**Figure 6j-l; Figure S8**).

## Discussion

*In vitro* myocardial models remain morphologically, molecularly, and functionally distinct from their *in vivo* counterparts, notwithstanding recent advancements in knowledge of CM maturation ^10,29,38^. Furthermore, the approaches used in culture medium design to date have been largely prescriptive, and factor selection has been limited by the feasibility of assaying quantitative and high-throughput markers of maturation.

Traditional regression-based optimization methodologies must be limited in scope to be logistically feasible and of sufficient statistical resolution to be actionable. We leveraged a directed evolutionary algorithm to survey a much wider set of additives over successive iterations, using a metric of metabolic organization and anabolism (UCR) that by self-normalization allowed for maximizing separate runs per generation. A range of UCR performance was observed across the multiple generations of iteration, but the final C16 formulation was validated to outperform gold-standard commercial and homemade media in all metrics tested, both quantitative and qualitative, making it a key candidate for further benchmarking. C16-treated iPSC-CMs displayed marked morphological changes, including hypertrophy, elongation, and sarcomeric striation evident under brightfield examination, and corresponding development of T-tubule-like (DHPR positive) and SR-like (SERCA2a and RyR2 positive) structures (**Figure 1**). Differential contractility was evoked by C16 treatment across two healthy iPSC-CM lines, as well as a CRISPR-induced mutant line.

The appearance of inward rectifier K^+^ currents (I_K1_) is a key milestone in our CM maturation approach, as it is needed to prevent spontaneous AP firing in working atrial and ventricular CMs ^39,40^. This quiescence is essential *in vivo* to allow nodal control of ventricular contraction combined with proper ordering of the ventricular contraction by Purkinje fibres, thereby ensuring efficient pumping action. C16-treated iPSC-CMs exhibited several markers of advanced electrophysiological maturation (**Figure 2**), including characteristic I_K1_ current ^35,39,41^. It is notable that C16-treated cells achieved I_K1_ densities roughly equivalent to those measured in isolated adult cardiomyocytes, although I_K1_ densities are known to display regional heterogeneity ^42^. Accordingly, C16-treated iPSC-CMs were generally quiescent after approximately 3 weeks, and did not display Aps or contractile spontaneity at 6 weeks unless I_K1_ was blocked by targeted Ba^2+^ application.

In addition to their electrophysiology, C16-matured iPSC-CMs demonstrated advanced Ca^2+^ handling and metabolism (**Figure 3**). Kinetics of steady-state paced Ca^2+^ transients were generally accelerated, and cells exhibited a marked kinetic response to isoproterenol stimulation, which is consistent with the observation of increased β-adrenergic response in maturing CMs ^6,13,43^. Importantly, this response relies not only on increased expression of the βadrenergic receptor, but also on the formation of functional dyads to potentiate SR^2+^ Ca store release. The presence of functional SR Ca^2+^ reuptake, as opposed to only sarcolemmal Ca^2+^ efflux, would also be consistent with the accelerated early decay kinetic parameters observed in baseline Ca^2+^ transients of C16-treated cells. Furthermore, upon challenge, matured cells demonstrated increased verapamil (an L-type current blocker) sensitivity and had significantly increased caffeine-sensitive SR Ca^2+^ stores. This recapitulation of advanced CM functionality was reflected in metabolic analysis, where cells demonstrated higher per-cell oxidative rates, as well as higher central carbon flux and a lower reliance on anaerobic glycolysis, all hallmarks of mature CM metabolism ^44^.

Finally, transcriptomic analysis of matured iPSC-CM monolayers (**Figure 4**) suggested strong evidence of functional maturation, including downregulation of products associated with differentiation, proliferation, and motility, and upregulation of products associated with excitation-contraction coupling, electrophysiology, lipid transport, and oxidative metabolism. Proteomic analysis (**Figure 5**) reflected many of the expressional changes expected to accompany increased contractility, improved Ca^2+^ handling, and metabolic function, including cytoskeletal, sarcoplasmic and trafficking, and substrate trafficking and OXPHOS ontologies, respectively. Targeted proteomic approaches may produce additional insight with respect to the electrophysiological profile of maturing iPSC-CMs. Furthermore, the transcriptional downregulation of key traditional markers of iPSC-CM maturity such as *MYH7* and *TNNI3* in the C16 treatment was not reflected in the expression of their protein products, suggesting that transcriptional assessment without corresponding protein or functional verification may not be sufficient in determining the maturation status of advanced iPSC-CM cultures.

This increase in hallmark CM function was also observed in Biowire II engineered myocardial tissues, where treatment with C16 medium was additive to continuous electrical field stimulation in evoking increased functional performance (**Figure 6**). Tissues treated with C16 medium and subjected to non-progressive electrical stimulation demonstrated steady-state twitch stresses near the peak of engineered myocardium at physiological [Ca^2+^] and at an appreciable fraction (*e*.*g*. ∼2-5%) of *ex vivo* measurements of healthy adult myocardium ^45–47^. Maturing function was further reflected in the highly positive force-frequency (Bowditch) response of C16-treated tissues, and in the tissue architecture, functional protein expression, and TEM-visible ultrastructural associated with myocardial maturation. Interestingly, the well-validated importance of electrical stimulation was replicated in our study, but overshadowed by the statistical effect of culture medium treatment in both functional and transcriptomic analysis. However, despite the transcriptomic similarity between unstimulated and stimulated C16-treated tissues, stimulated tissues consistently outperformed the unstimulated condition on physiological analyses, suggesting non-transcriptomic contributions to tissue function. Efforts to further improve tissue functionality could include extended culture periods, the use of progressive stimulation protocols ^24,43^, or increasing cell diversity in co-cultures.

Several previous studies have examined the transcriptomes of maturing PSC-CMs ^13,35,43,48–51^ or native myocardium ^13,52–54^, but the majority of maturation protocols have been characterized only relative to freshly-differentiated PSC-CMs or time-matched gold standards. Using these internal controls, we were able to construct a meta-analysis of CM maturation both *in vitro* and *in vivo*. The positioning of the maturation protocol clusters relative to the *in vivo* clusters appeared to indicate that 2D approaches promote maturation of the transcriptome most effectively, while EB approaches reduce maturity relative to controls. Analysis of the PC loadings challenges this interpretation, as functional categories associated with CM maturation could be identified in all PC loadings diverging from freshly-differentiated PSC-CM controls. Furthermore, each maturation protocol included in the meta-analysis has been experimentally shown to promote functional maturation in some capacity, even in EB methods which most oppose *in vivo* samples in the PCA. However, in this study and others, 3D co-culture of CMs and fibroblasts more closely replicates the mechanical and biochemical niche of myocardium and evokes emergent functional physiological characteristics and metrics that have not to date been achieved in simple monolayers; their transcriptomic trends are further consistent with marked downregulation of proliferative ontologies, but incomplete downregulation of developmental gene programs. Collectively with our insights from paired proteomic experiments, these observations therefore suggest that functional maturation may not be required to follow the same linear trajectory observed for the transition from fetal to adult CMs in vivo, and that achieving an ‘adult-like’ transcriptional profile may not be absolutely required for functional maturation. Additionally, the direct interpretation of *in vitro* transcriptomic datasets in the context of *in vivo* samples is complicated by the wealth of cell types in myocardial tissue, including endothelial cells, fibroblasts, pericytes, and immune cells ^55^, which even if removed from digested samples bound for sequencing, may exert regulatory control on CM phenotype *in situ*.

The development of a highly-mature PSC-CM source will enable critical advancements in several downstream applications, particularly ones that require PSC-CMs at an advanced stage of electrophysiological development. Cell transplantation applications of PSC-CM technology have been complicated by the presence of graft-induced focal arrhythmias, ostensibly due to continued AP spontaneity ^56^. Furthermore, Torsades de Pointes ventricular tachyarrhythmias result from both genetic and pharmacological risk and are notoriously difficult to model *in vitro*; TdP stems from hERG (human ether-a-go-go; K_v_11.1) current blockade, but this current is highly dependent on other coexisting CM currents, complicating pharmaceutical design and predictive preclinical evaluation ^8,9^. Further development and in-depth examination of high-fidelity and electrophysiologically mature iPSC-CMs may provide insight on their suitability for cardiotoxicity screening applications.

In conclusion, the iteratively-optimized C16 medium significantly advanced maturation of PSC-CMs toward that of mature adult tissues in both monolayer and co-culture organoids; this formulation may be beneficial for applications in basic physiology, drug screening, personalized medicine, or the design and execution of cell or tissue therapies. This study also directly compares a large collection of CM maturation datasets within a single analysis, providing insight into potential lacking physiological ontologies in existing maturation protocols (including the one presented in this study). Furthermore, within our own study, we demonstrate inconstancies between transcriptomic, proteomic, and functional analyses, demonstrating the importance of the use of different levels of assessment of physiological maturity. There is likely significant benefit to further testing of novel iPSC-CM maturation medium formulations, with additional additives that may either push maturation signaling or supplement crucial metabolic processes. By providing a framework to achieve advanced levels of hallmark functional iPSC-CM metrics, our findings suggest that the process and components underlying functional CM maturation may be more highly regulated and multifaceted than previously understood, and provide a basis for the generation of novel hypotheses regarding the process and regulation of CM maturation. More generally, this study provides a proof of concept for high-dimensional optimization of stem cell-derived cultures and engineered tissues.

## Supporting information

Proteomics sample IDs

Video S1: C16-treated unstimulated Biowire

Video S2: RPMI + B27-treated unstimulated Biowire

Video S3: C16-treated stimulated Biowire

Video S4: RPMI + B27-treated stimulated Biowire

## Acknowledgements

The authors acknowledge Dr. Michael Laflamme (University Health Network) for constructive conversations in the ideation and realization of this project. The authors also acknowledge The Metabolomics Innovation Centre (TMIC; McGill University Node, Montreal QC), the Nanoscale Biological Imaging Facility (NBIF) at the Hospital for Sick Children (Toronto ON), and the Princess Margaret Genomics Centre (PMGC, University Health Network, Toronto ON) for their assistance with metabolomics, transmission electron microscopy, and RNA sequencing, respectively. This study was funded by a Canadian Institutes of Health Research (CIHR) Project grant (PJT-175231) to CAS; a Collaborative Health Research Program grant from CIHR (CPG-151946) and the Natural Sciences and Engineering Research Council of Canada (NSERC) (CHRPJ 508366-17) to CAS and FB; a Ted Rogers Centre for Heart Research Strategic Innovation Grant to JE, SM, FB, MR, and CAS; a Canada Research Chair in Stem Cell Models of Childhood Disease to JE; and a Heart and Stroke Foundation of Canada / Robert M. Freedom Chair in Cardiovascular Science to SM. NIC and RGI were funded by Vanier Canada Graduate Scholarships, from NSERC and CIHR, respectively. LJD was funded by the Translational Biology and Engineering Program, Ted Rogers Centre for Heart Research. This paper was typeset with the bioRxiv word template by @Chrelli: www.github.com/chrelli/bioRxiv-word-template.

## Author contributions

NIC and CAS conceived the study. NIC, MMK, and JA contributed to the iterative optimization workflow. NIC, LJD, WC, UK, MZM, ZM, and RGI collected and analyzed data. JPS, AOG, SM, JE, PHB, and CAS supervised the study. NIC, LJD, and WC prepared display items and drafted the manuscript. All authors edited the manuscript and approved the final version.

## Competing interest statement

NIC and CAS have filed a provisional patent in partnership with the University of Toronto for the formulation described in this study. No other authors claim a conflict of interest.

## Methods

No statistical methods were used to predetermine sample size. Assignment of experimental wells was randomized, but blinding of conditions could not be used during experiments or outcome assessment due to physical differences (*e*.*g*., medium colour) between the treatments used and the morphological differences in the resulting cells.

### Design of an iterative algorithmic search for optimized maturation medium

The C16 maturation medium was non-prescriptively optimized in a large solution space (∼763 billion discrete formulations for an analogous 5^17^ full factorial experimental design), with generations of formulations iteratively scored and evaluated according to a novel objective metric, the respirometric uncoupled:control ratio (UCR) as measured by Seahorse XFe96 analysis. This design philosophy allowed for agnostic testing and the evolution of emergent physiological interactions between multiple defined soluble factors within a formulation; several of these factors had not been surveyed previously for efficacy toward PSC-CM maturation. Additionally, we used exclusively M199-based formulations for two reasons: firstly, the ionic profile of M199 is closer to that of human plasma than DMEMor RPMI 1640-based media, especially in [Ca^2+^], for which homeostasis is vital to developing CMs; and secondly, that M199 contains a much more diverse source of cofactors and vitamins than either basal DMEM or RPMI 1640 media. For the latter consideration, the use of a defined formulation negates the possibility of serum providing these essential components, especially in a highly-specialized cell such as a maturing CM, which may outsource many non-core biosynthetic and secondary metabolic pathways to other tissues of the body.

The formulation search was performed using an existing iterative High-Dimensional, Differential Evolution (HD-DE) decision tree algorithm programmed in MATLAB ^32^; the published script was modified for R2018a version compatibility, for the use of 17 variable factors, and for duplicate runs. Briefly, the algorithm generates formulation candidate vectors within a defined number of runs, factors, and levels of said factors. Maximal coverage during the initial generation is generated using a Sobol quasi-random distribution (e.g., minimizing discrepancy). After a generation is produced, objective metric scores are reported, theoretically forming a rough Pareto distribution. Formulations within 10% of the top scorer are chosen to continue in the next generation, while poor-scoring formulations are either replaced with randomly-generated vectors, or with offspring vectors from the intergenerational memory of top scorers. The process is formally terminated when the median candidate of the Pareto-ranked generation does not improve by at least 10% after 3 consecutive generations.

### Materials

For iterative formulation candidate screening and characterization of the C16 formulation, soluble components used were 2-deoxyglucose (DXG498; BioShop), sodium L-lactate (L7022; Sigma), Chemically Defined Lipid (11905031; Gibco), D-galactose (GAL500; BioShop), creatine monohydrate (CREE200; BioShop), L-carnitine hydrochloride (C0283; Sigma), taurine (TAU303; BioShop), insulin-transferrin-selenium (41400045; Gibco), bovine serum albumin fraction V (10735078001; Roche), β-mercaptoethanol (M3148; Sigma), putrescine dihydrochloride (PUT001; BioShop), ethanolamine hydrochloride (E6133; Sigma), triiodothyronine (T6397; Sigma), recombinant insulin-like growth factor 1 (PHG0071; Gibco), recombinant leukemia inhibitory factor (SRP3316; Sigma), growth hormone (869008; Millipore Sigma), hydrocortisone (H0888, Sigma), recombinant neuregulin 1β2 (ab73753; Abcam), metformin hydrochloride (ICN15169101; MP Biomedicals), in a base of Medium 199 (M4350; Sigma). Gold-standard controls used were iCell cardiomyocytes maintenance medium (CMM-100120-001; Cellular Dynamics), cardiomyocyte maintenance medium (05020; STEMCELL Technologies), and RPMI 1640 (R8758; Sigma) with 1X B27 supplement (17504044; Gibco). An additional top-performing formulation from recent literature (“Feyen”) was prepared as previously described ^35^ and used in proteomic and metabolomic analyses as an exemplar of maturation progress.

For immunofluorescent staining, anti-α-actinin (ab9465; Abcam) and anti-connexin 43 (ab217676; Abcam) were used, as well as phalloidin-tetramethylrhodamine B isothiocyanate, (P1951; Sigma), and DAPI (62248; Thermo Scientific). For respirometry, a DMEM-based XF medium (103334; Agilent) was used, with oligomycin (O4876; Sigma), FCCP (15218; Cayman Chemical), antimycin A (A8674; Sigma), and rotenone (NC0779735; Cayman Chemical) used as inhibitors. Unless otherwise noted, all other materials were obtained from Sigma.

### Cells

An iterative formulation search, and subsequent characterization of iPSC-CM cultured in the chosen final maturation medium candidate (“C16”) was performed using the Personal Genome Project of Canada-17 iPSC line (PGPC17), which have been extensively characterized across a wide range of differentiation pathways ^33^. Further testing for cross-cell line compatibility was performed on PGPC14 iPSCs, and on *MYBPC3*-knockout PGPC17 cells generated via CRISPR; both lines were also previously characterized as iPSC-CMs ^33^.

Cells were frozen in clumps in mTeSR (STEMCELL Technologies) containing 10% DMSO, and thawed and cultured when ready for differentiation. Cells were cultured in mTeSR on Geltrex (Thermo-Fisher; LDEV-free, hESC-qualified; incubated at 1:100 in DMEM for 1 h on TCPS plates) and passaged in clumps using ReLeSR (STEMCELL Technologies), all according to manufacturer guidelines. Cells were passaged at least two times after thawing before being expanded for differentiation. PSC-CM differentiation was performed using the Cardiomyocyte Differentiation Kit (STEMCELL Technologies) according to manufacturer directions, with cell seeding densities optimized for each cell line (between 4-8 × 10^5^ cells well^−1^ in a 12well dish, previously coated with 1:100 Matrigel (Corning) in DMEM). Cells were cultured post-differentiation until d18 post induction before reseeding for maturation experiments. Obtained monolayers in 12-well format were dissociated within 1 week of differentiation by incubation for 1 h at 37°C in 1 mL Hanks buffer, containing 200 U mL^−1^ Collagenase Type II (Worthington), with 0.5 mL TryPLE Select (Gibco) added for 15 additional minutes. Cells were centrifuged 5 min at 300 x *g*, resuspended, and plated in RPMI + B27 for 1 d culture before applying treatments. Non-screening treatments were applied to monolayers for 6 weeks; all monolayer cultures were performed in the absence of antibiotics.

### Formulation screening

Soluble factors used in the iterative screening and final C16 formulation (**Table S1**) were selected with the goal of providing a range of substrates, cofactors, and hormones that would not necessarily be created by a maturing or mature CM but which would be conducive to increased biosynthesis and hypertrophy, oxidative metabolism to fuel highly-energetically demanding physiological processes, or activation of pathways implicated in maturation of myocardium or other functional tissues *in vivo*. In all, 169 unique formulations were queried over 4 generations of varying size, using the Seahorse XFe96 mitochondrial stress test to interrogate metabolic function.

Cells were dissociated from differentiation wells and seeded at 8 × 10^4^ cells well^−1^ in CM support medium on Matrigel-coated XFe96 monolayer plates (Agilent Technologies). Culture medium was changed the next day to the well’s respective treatment. Cells were then cultured 21 days in 60 μL medium well^−1^, with full medium changes every second day for 3 weeks in either duplicates of a formulation candidate, or one of four wells of goldstandard STEMCELL Technologies Cardiomyocyte Maintenance Medium. The four corner wells of the plate were additionally left unseeded as internal measurement controls according to standard XFe96 manufacturer guidelines.

### Respirometry

Cells were equilibrated 45 min before metabolic characterization in initial volume 150 μL Seahorse XF base medium (103334-100, Agilent Technologies) containing additional (in mmol L^−1^) glucose (5), pyruvate (1), glutamine (1), sodium lactate (5), and 1X Chemically Defined Lipid, to best supply the oxidative substrate flexibility of mature CMs ^75^. A mitochondrial stress test was performed with sequential injections of 25 μL each (final concentrations): oligomycin (2.5 μmol L^−1^), carbonyl cyanide-4-phenylhydrazone (FCCP; 1 μmol L^−1^ for iterative testing, or either 0.20, 0.40, 0.50, 0.65, 0.80, or 1.00 μmol L^−1^ for characterization of iPSC-CMs cultured in C16 medium), antimycin A and rotenone (2.5 μmol L^−1^ each). Due to the high specific oxidative flux of CMs, a modified measurement protocol to avoid hypoxia was employed ^76^; cells were measured for 2 rather than 3 cycles at each step, with the minimum measurement time of 2 min.

Respirometric measurements were normalized to cell numbers per well. Each well was fixed in 2% formalin for 5 min, washed 3 times with Ca^2+^ and Mg^2+^-free PBS, stained with Hoechst (1 μg mL^−1^) in PBS without for 5 min, washed 3 additional times, and imaged using an IX71 inverted widefield fluorescent microscope (Olympus Corporation) with a FITC filter. Cell quantifications were performed by performing an automated count of nuclei using ImageJ 1.52p (NIH), by sequential use of the standard Otsu threshold, watershed segmentation, and particle count functions. Normalized per-cell oxidation rates (OCR) and uncoupled:control ratios (UCR) were compared between treatments using a repeated measures ANOVA and Dunnett’s multiple comparisons post-hoc test for gold-standard treatments vs. the C16 formulation where indicated, to account for batch effects of the respirometric assay.

### Immunofluorescence and confocal microscopy

Cultured cells were fixed with 2% paraformaldehyde for 10 min at room temperature, followed by 90% ice-cold methanol for 10 min, followed by permeabilization buffer (0.5% Triton X-100, 0.2% Tween-20 in PBS) for 30 min at 4 °C. Blocking buffer (5% FBS in permeabilization buffer) was then added and incubated for 1 h at room temperature. Cells were incubated with primary antibodies (α-actinin 1:500, Cx43 1:500) in blocking buffer overnight at 4 °C; incubation in rhodamine-red-conjugated phalloidin (Sigma) was 1 h at RT. Fluorophore-conjugated secondary antibody staining (anti-rabbit AlexaFluor® 488, anti-mouse AlexaFluor® 594, anti-mouse AlexaFluor® 647; Molecular Probes; 1:800) was performed at room temperature for 1 h in the dark. Nuclear counterstaining was performed using 1 μg ml^−1^ DAPI at room temperature for 15 min in the dark. Images were acquired using an FV3000 confocal microscope with 405, 488, 561, and 640 nm lasers.

### Optical contractile tracking of iPSC-CM monolayers

Contractile kinetics were assessed from phase-contrast videos recorded at ∼20 fps with a 40X objective, using a particle-image velocimetry package for ImageJ ^77^ to quantify relative displacement from the relaxed state between contractions. Individual traces from biological replicates were fitted with a quartic regression using Prism 9 (GraphPad Software Inc., San Diego, USA).

### Electrophysiological characterization

#### Intracellular recordings

Action potentials (APs) were record in single iPSC-CMs or small clusters of iPSC-CMs (n = 14 for C16 and n = 30 for RPMI + B27) using sharp microelectrodes of resistances between 40 to 90 MΩ (filled with 3 mol L^−1^ KCl) with an Axopatch 200B amplifier. Intracellular voltages were digitized at a sample rate of 10 kHz (Digidata 1440 A plus pCLAMP 10.3, Molecular Devices). Voltage recording was performed at room temperature and perfused with modified Tyrode’s solution (pH 7.4) containing (in mmol L^−1^): NaCl (120), KCl (5) CaCl_2_ (2), MgCl_2_ (1), Na_2_HPO_4_ (0.84), MgSO_4_ (0.28), KH_2_PO_4_ (0.22), NaHCO_3_ (27), and glucose (5.6) ^39^.

#### Patch-clamp recordings

Voltage-clamp measurements were used to record whole-cell currents in single isolated iPSC-CMs using the patch-clamp technique at room temperature. The hardware was the same as for the AP recordings. The pipette solution (pH 7.2 adjusted with KOH) contained (in mmol L^−1^): K-gluconate (150), EGTA (5), HEPES (10), and MgATP (5). The series resistance of the pipettes was 3-4 MΩ. The external bath solution (pH 7.35-7.4 with NaOH) contained (in mmol L^−1^): NaCl (148), KCl (5.4), MgCl_2_ (1.0), CaCl_2_ (1.8), NaH_2_PO_4_ (0.4), glucose (5.5), and HEPES (15). The external perfusion solutions also contained nifedipine (5 μmol L^−1^) to block possible overlapping Ca^2+^ currents. I_K1_ was assessed by using voltage-steps from −115 mV to −20 mV (in 5 mV intervals) were applied for 500 ms from a membrane voltage of −60 mV. Currents were recorded at sampling rates of 10 kHz and analyzed using Clampfit 10.7 (Molecular Devices). Current-voltage (I-V) relationships were generated using peak currents as a function of membrane voltage. Currents were also recorded following the perfusion of BaCl_2_ (100-400 μmol L^−1^), which is a potent and selective blocker of I_K1_ at such low concentrations ^78^.

#### Ca^2+^ imaging

Cells cultured in 12-well plate format were dissociated as described above and replated in single-cell format on 96-well plates with a #1.5 coverslip bottom treated with 1:100 Matrigel, and cultured in their treatment medium to recover for 2 days before imaging. Frozen dry vials of 50 μg Fluo4 AM (Thermo-Fisher) were reconstituted with 50 μL DMSO containing 20% (w/v) Pluronic F-127. Cells were treated with 50 μL well^−1^ Fluo-4 AM solution, diluted 1:100 in Tyrode’s solution, for 30 min at RT. Cells were rinsed twice with Tyrode’s solution and maintained up to 2 h thereafter in 100 μL Tyrode’s solution, containing 5 μmol L^−1^ S-blebbistatin (Toronto Research Chemicals) to prevent movement artifact.

Ca^2+^ imaging was performed on a FluoView FV3000 (FV3000; Olympus Corporation) confocal microscope heated to 37 °C. Linescan measurements with 488 nm excitation and collection from 505-550 nm at 1% laser power and c.a. 500 V at the HSD were taken at the 20X objective and 2X zoom with spatial and temporal resolutions of 0.62 μm and c.a. 1.2-1.8 ms, respectively. Cells were subjected to steady-state stimulation at 1 Hz monophasic square-wave impulses of 2 ms and 40 V at approximately 0.6 cm using custom-made graphite electrodes connected to an S48 physiological stimulator (Grass Technologies, Warwick, RI). For monolayer SR capacity experiments, cells were paced at 1 Hz at steady-state before consecutive additions of verapamil (20 μmol L^−1^) and caffeine (10 mmol L^−1^). Fractional SR load upon addition of caffeine was normalized to steady-state transient amplitudes for each cell; comparisons did not assume equal variance and were analyzed by Welch’s t-test. Fluo-4 fluorescence measurements were taken using a CCD camera at c.a. 30 Hz framerate on an IX71 inverted epifluorescence microscope (Olympus Corporation, Tokyo, Japan) equipped with a FITC filter cube. Transients were manually extracted using ImageJ. Epifluorescent SR capacity transients were directly reported and interpreted. Transient confocal linescans were smoothed with a 7-frame rolling-average filter and analyzed for kinetics, using a custom MATLAB script. Kinetics parameters assessed included time from baseline to peak fluorescence, and times from peak fluorescence to a decay of 10%, 25%, 50%, and 75% peak magnitudes. Kinetic metrics were compared between treatments using a Brown-Forsythe ANOVA and Dunnett’s multiple comparisons post-hoc test for gold-standard treatments vs. the C16 formulation where indicated, to account for unequal variances between treatments.

### Microtissue maturation

#### Culture and functional benchmarking

Microtissues were created according to established methods ^36^. Briefly, iPSC-CMs mixed with primary human fibroblasts were cast in fibrin gels within embossed polystyrene molds and suspended between flexible poly(octamethylene maleate (anhydride) citrate) (POMaC) posts to allow for contraction against resistance. All strips were cultured in 10 cm cell culture dishes, containing 10 mL of medium changed twice weekly, with 0.1% penicillin-streptomycin. The C16 medium used for monolayer culture was found to be incompatible with tissue function due to toxicity to primary cardiac fibroblasts (as found in monolayer fibroblast culture); a modified version without 2-deoxyglucose allowed continued tissue function and was used as the working version of C16 for all Biowire II experiments. Strips of up to 8 functional tissues were cultured 1 week in RPMI+B27 medium, to allow compaction, before assignments to treatments in either modified C16 2-deoxyglucose was omitted for co-culture applications, as it was found to be toxic to primary fibroblasts at working concentrations) or continued RPMI + B27 media, and with or without electrical field stimulation (10 V cm^−1^ at 1 cm, 1 Hz monophasic square wave pulses for 3 ms) via two graphite electrodes connected to an S48 physiological stimulator (Grass Technologies, USA). Tissues were assessed for function at 0, 7, 14, and 21 d of treatment for baseline paced diastolic stress, peak systolic twitch stress, and stimulation force-frequency relationship up to 6 Hz using an IX71 microscope at 37 °C. Polymeric wires were separately assessed for stiffness using a MicroSquisher (CellScale, Waterloo, Canada) to calculate absolute tissue diastolic and active forces. Twitch forces were calculated as the difference between diastolic and active forces. Stresses were derived from forces normalized to calculated cross-sectional tissue area (an ellipse of 5:3 aspect ratio at the tissue midpoint between polymeric wires based on existing cross-sectional imaging ^36^). Stress values were first analyzed using a 3-way repeated measures ANOVA (Medium x Stimulation x Time) to assess interactions between the effects of medium versus stimulation on systolic stress. If the 3-way interaction analyses failed to find significance, the were repeated by 2-way repeated measures ANOVA combined with a within-timepoints Tukey’s HSD, as post-hoc test. Sphericity was not assumed and the Geisser-Greenhouse correction was applied when indicated.

### Transmission electron microscopy

Microtissues were rinsed 3x with PBS before fixation in 4% paraformaldehyde and 1% glutaraldehyde in PBS. Secondary fixation was in 1% OsO_4_ in 0.1 mol L^−1^ sodium cacodylate buffer. Specimens were dried in a series of 50%, 70%, 90%, and 3x 100% EtOH, then infiltrated in 50% and 70% (2 h steps) and 100% (overnight) Spurr resin. Samples were sectioned at 70 nm, stained with uranyl acetate and lead citrate, and imaged with a T20 transmission electron microscope (Thermo) and a 4k CCD (Gatan).

### ^13^C metabolic flux analysis

Cells were incubated 24 h in one of three substrate-labeled culture media. The base medium consisted of glucose-free DMEM, containing 0.1% w/v Fraction V bovine serum albumin, 50 μmol L^−1^ carnitine, and 1X InsulinTransferrin-Selenium supplement (Gibco). Medium also contained 10 mmol L^−1^ glucose, 3 mmol L^−1^ lactate, and 120 μmol L^−1^ palmitate; in any well, one of these three carbon sources was U-^13^C labeled while the remaining two were untagged. After incubation, cells were quickly washed twice in PBS and extracted with a 2:2:1 (v:v:v) mixture of acetonitrile, methanol, and water, all mass spectrometry grade and pre-cooled to −80°C. Cells were scraped from plates and the slurry kept at −80°C until analysis. Analysis was performed by LC-MS by The Metabolomics Innovation Centre at McGill University according to previously-established methods ^79^, and isotopomer relative balances were measured for metabolites of interest with each label.

### Protein extraction and global proteomic profiling

Cells were washed 3x in PBS, scraped from 12-well plates, and centrifuged 5 min at 200 x *g*. Pellets were flash-frozen and stored at −80°C until extraction. Protein extraction was in 8 M urea with 5×3 s sonication bursts at 30% power on ice. Protein concentrations were assessed by BCA assay (Pierce). Samples were reduced in 2.5 mmol L^−1^ dithiothreitol (DTT) 1 h at 37°C, before alkylation in 5 mmol L^−1^ iodoacetamide for 30 min at RT in the dark. Finally, samples were diluted 1:10 in 50 mM ammonium bicarbonate before the addition of 1:50 protein:protein by mass of sequencing-grade trypsin (Promega) for overnight digestion at 37 °C, followed by the terminating addition of formic acid at a final concentration of 1%. Mass spectrometry and analysis of proteomic samples were performed as previously described ^80^. Mass spectrometry proteomics data have been deposited to the ProteomeXchange Consortium via the PRIDE partner repository with the dataset identifier PXD036639.

### RNA sequencing

#### Sample Preparation

Total RNA was extracted from iPSC-CM monolayers using a Qiagen RNeasy Micro kit and prepared for sequencing at 100 ng with a Stranded Total RNA Prep kit (Illumina Inc., USA), and bulk-sequenced at the Princess Margaret Genomics Center (Toronto, Ontario) using 100 bp paired-end reads at a depth of 40 million reads. Biowire II tissues were extracted using a PicoPure™ kit (Thermo-Fisher), prepared with a SMARTer Stranded Total RNA-Seq Kit v3 – Pico Input Mammalian kit (Takara Bio Inc., Japan) with 0.75 ng and bulk-sequenced using 100 bp paired-end reads at a depth of 50 million reads. Reads were aligned to the human reference transcriptome assembly (GR.Ch38) using Salmon ^81^. Raw and processed RNAseq data have been deposited in the NCBI Gene Expression Omnibus under the GEO Series accession number GSE214617.

#### Data Analysis

All RNAseq data analyses were performed in R. Gene-level read counts were obtained using the Tximeta ^82^ package. For the meta-analysis, ComBat-seq ^83^ was used to correct for batch effects introduced by the inclusion of samples from different publications. For principal component and T-distributed stochastic neighbor embedding (tSNE) analyses, the raw count matrix was adjusted using the variance stabilizing transformation implemented in the DESeq2 package. PCA and PC-associated GO term enrichment was performed using the pcaExplorer ^84^ package. tSNE analysis was performed using the Rtsne package, and clusters were determined from tSNE analysis using the Louvain method ^85^, implemented in the bluster package. Differential expression was determined using the DESeq2 ^86^ package, and Benjamin-Hochberg adjusted p-values < 0.05 were considered significant. Gene set enrichment analysis (GSEA) was performed using the GAGE package ^87^, and terms with Benjamini-Hochberg adjusted p-values < 0.05 were considered significant. GO term similarity was determined using the kappa similarity index, and clusters were identified by hierarchical clustering. Pathway enrichment analysis was performed using the GAGE package ^87^, and pathway diagrams were generated using Pathview ^88^. All plots were produced using the ggplot2 and ComplexHeatmap packages.

**Figure S1.**
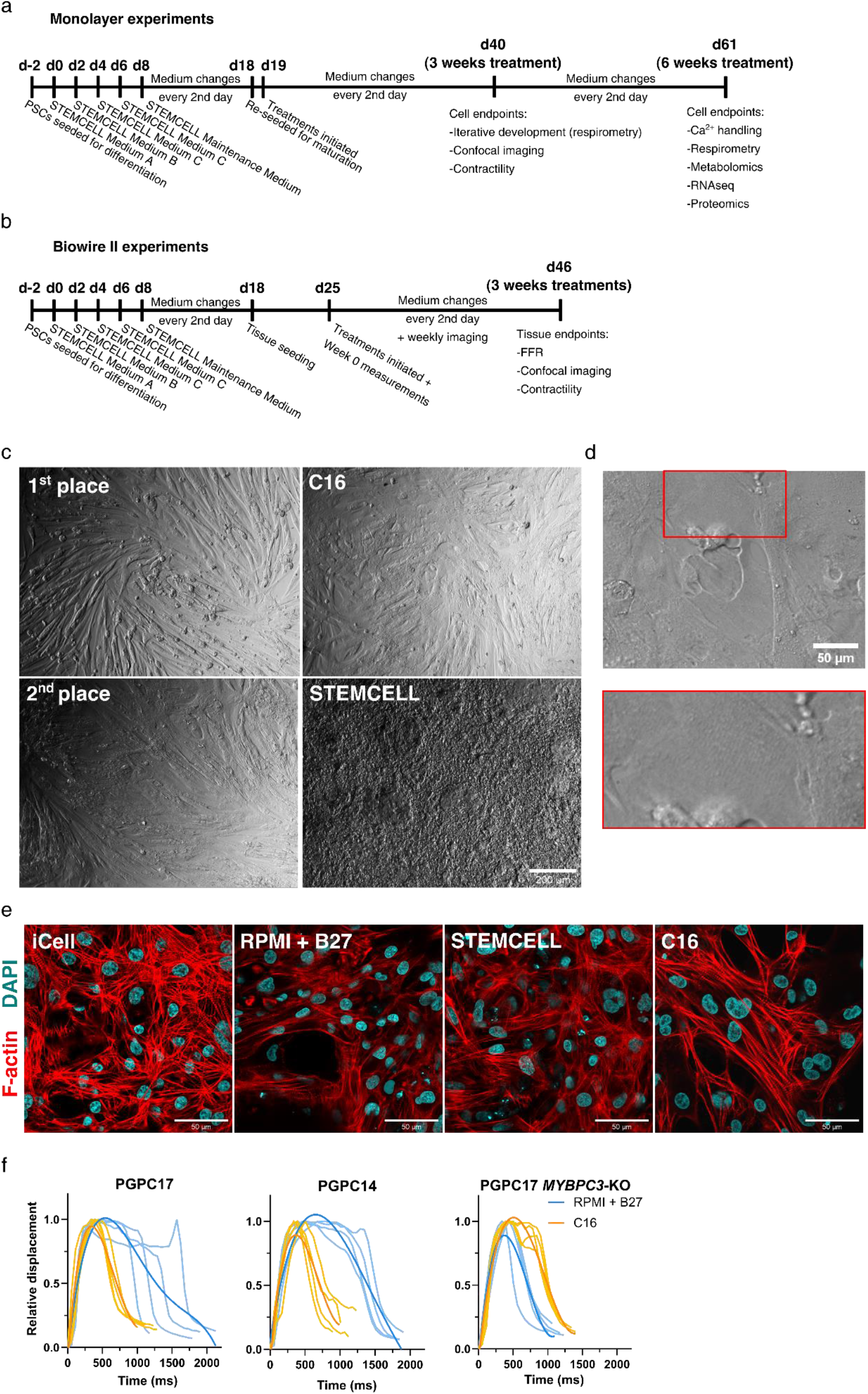
Experimental design, and morphology and performance of maturing iPSC-CM monolayers. **a)** Timeline of iPSC-CM differentiation, replating, monolayer treatment, and live analysis or harvesting for experiments described in the study. **b)** Timeline of iPSC-CM differentiation, Biowire II seeding, treatment, and live analysis or harvesting for experiments described in the study. **c)** Relief contrast microscopy reveals spontaneous elongation and local alignment of iPSC-CMs cultured 3 weeks in candidate maturation medium formulations (1^st^ and 2^nd^ place formulations by uncoupled:control ratio (UCR) metric in the iterative optimization process, and C16 medium) and STEMCELL Technologies Cardiomyocyte Maintenance Medium as a commercially-available gold-standard control. **d)** iPSC-CMs cultured 3 weeks in C16 medium have sarcomeric striations visible under brightfield microscopy; inset shows increased magnification to highlight striations. **e)** F-actin striation and bundling (red) visualized by phalloidin staining (DAPI in cyan) in cells cultured in C16 medium vs. iCell Cardiomyocytes Maintenance Medium, homemade RPMI + B27, or STEMCELL medium. **f)** Normalized displacement curves of spontaneously-contracting C16or RPMI + B27-treated iPSC-CM monolayers (individual traces pale, n = 4 each treatment; quartic fits dark) derived from PGPC17, PGPC14, and PGPC17-MYBPC3-KO iPSC lines. All analyses performed after 3 weeks of treatment with medium formulations.

**Figure S2.**
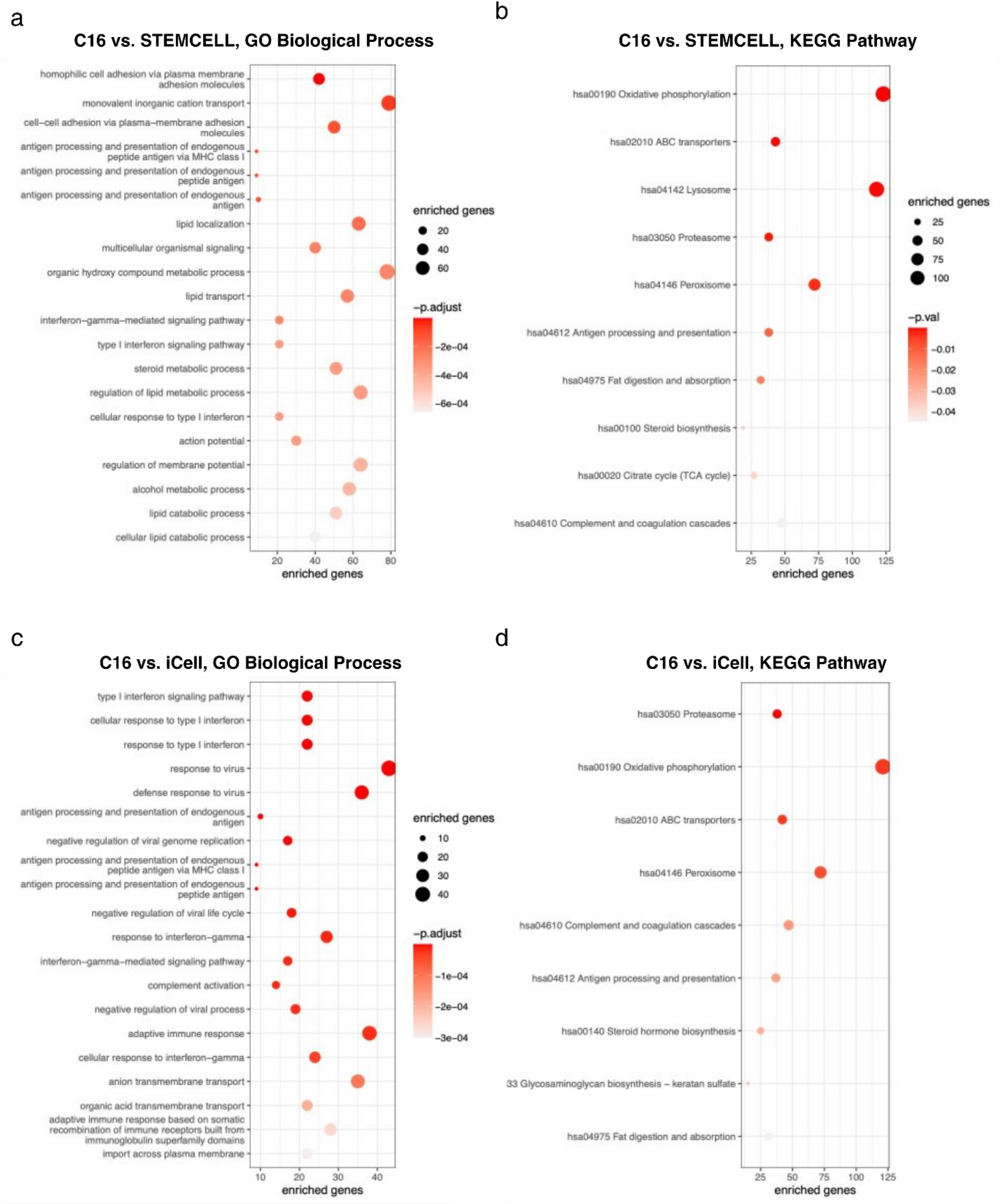
Transcriptionally-enriched functional terms and pathways in C16-treated iPSC-CMs. **a)** Top 20 enriched GO Biological Process terms among significantly upregulated genes in the indicated comparisons. Point size denotes the number of significantly upregulated genes in each enriched category. **b)** Top 20 enriched KEGG pathways in the indicated comparisons. Point size denotes the size of each gene set.

**Figure S3.**
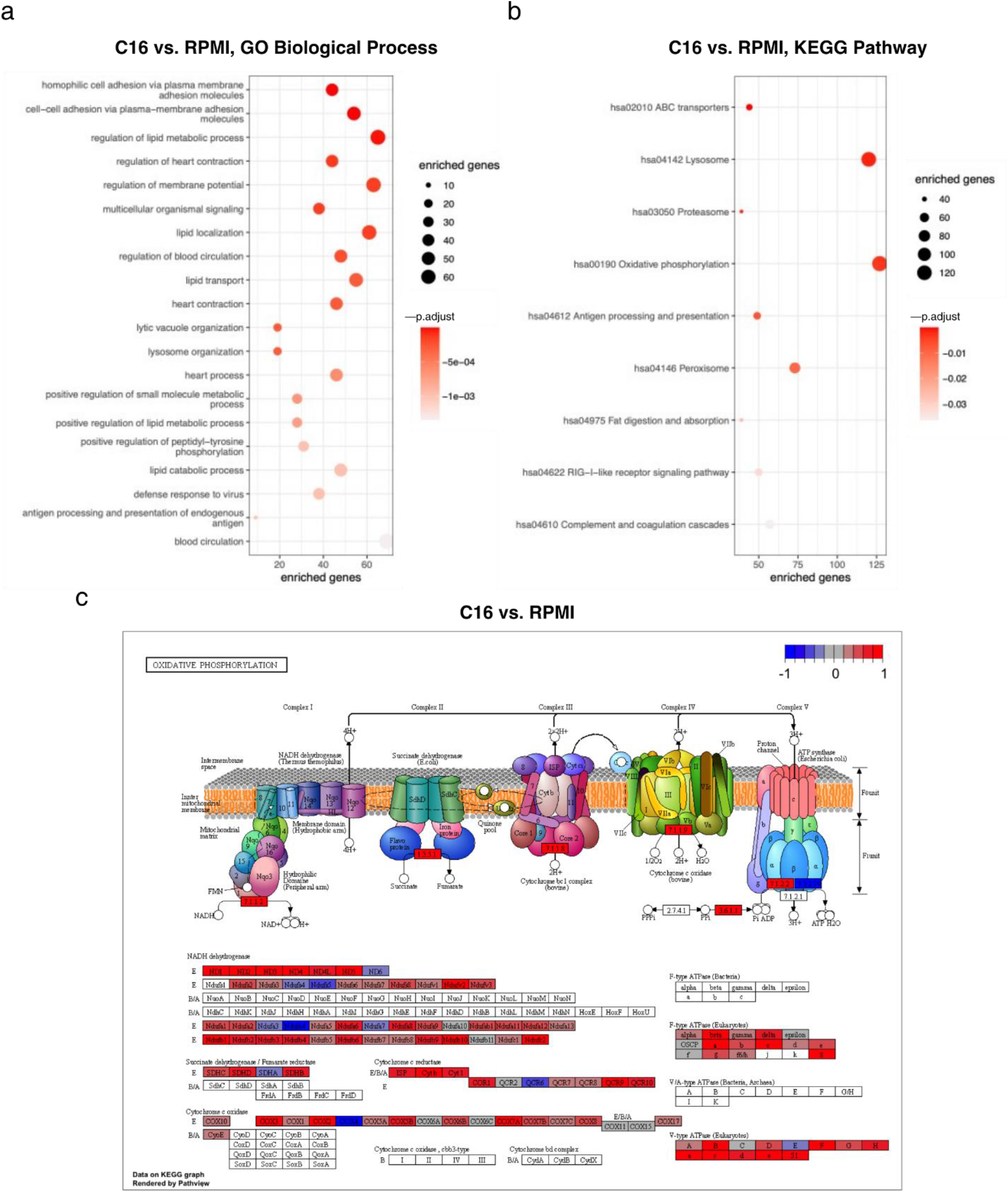
Transcriptionally-enriched functional terms and pathways in iPSC-CMs treated with C16 maturation medium. **a)** Top 20 enriched GO Biological Process terms among significantly upregulated genes in the indicated comparisons. Point size denotes the size of the enriched gene set corresponding to each term. **b)** Top 20 enriched KEGG pathways in the indicated comparisons. Point size denotes the size of the enriched gene set corresponding to each pathway. **c)** Schematic representation of the oxidative phosphorylation pathway (KEGG hsa000190) and corresponding gene expression data in C16 vs. treated iPSC-CMs. Color denotes log2 fold change relative to control.

**Figure S4.**
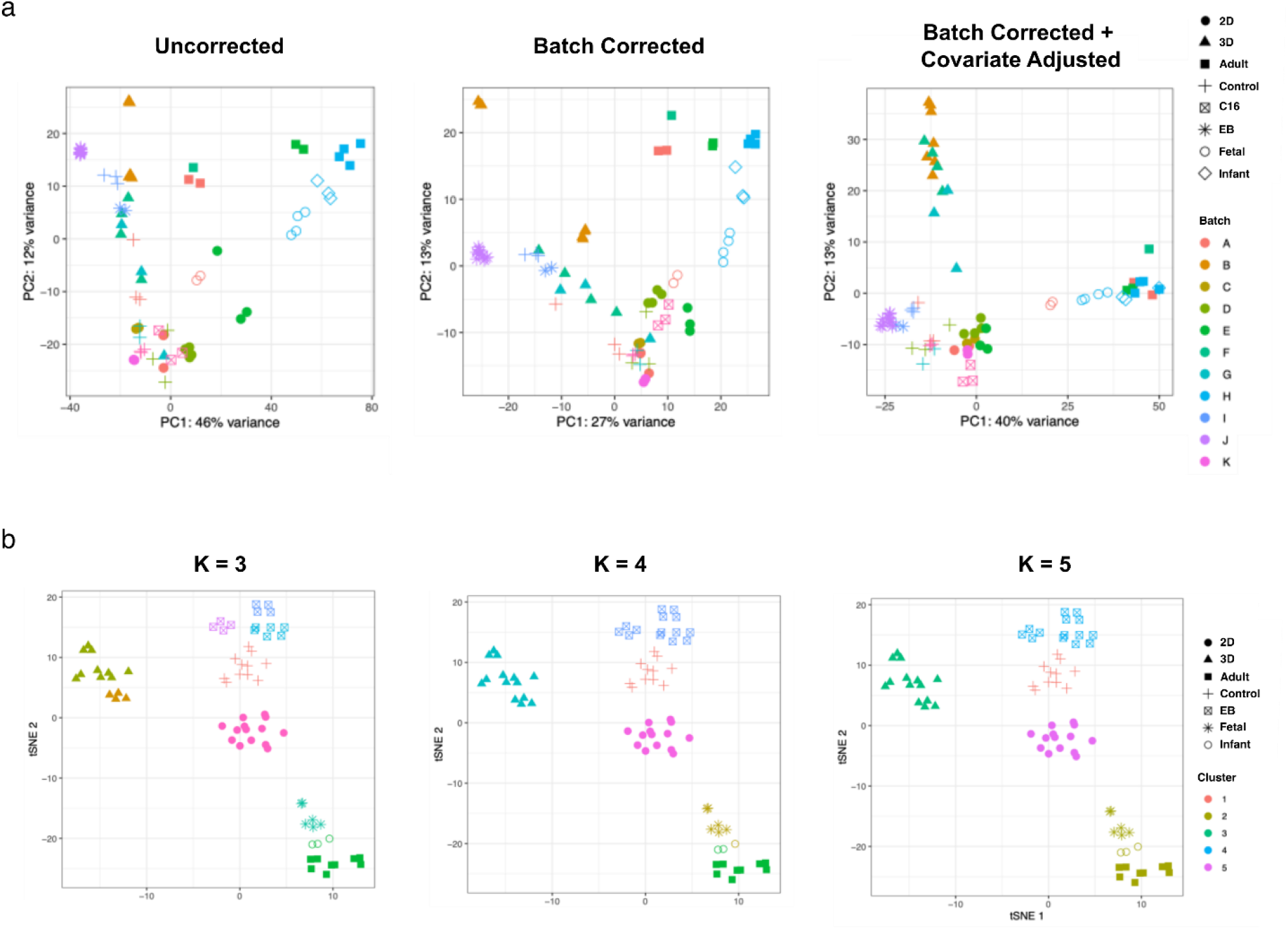
Validation of RNA-seq meta-analysis workflow. **a)** Principal component analysis of meta-analysis samples without batch correction (left), with batch correction only (middle), or with batch correction and specification of ‘type’ as a covariate parameter (right). Color denotes batch, and shape denotes sample type. See **Table S2** for assignment of samples to batches. **b)** tSNE plots displaying assignment of samples to clusters, using different k-values for construction of the k-nearest neighbour graph. Color denotes assigned cluster, and shape denotes sample type.

**Figure S5.**
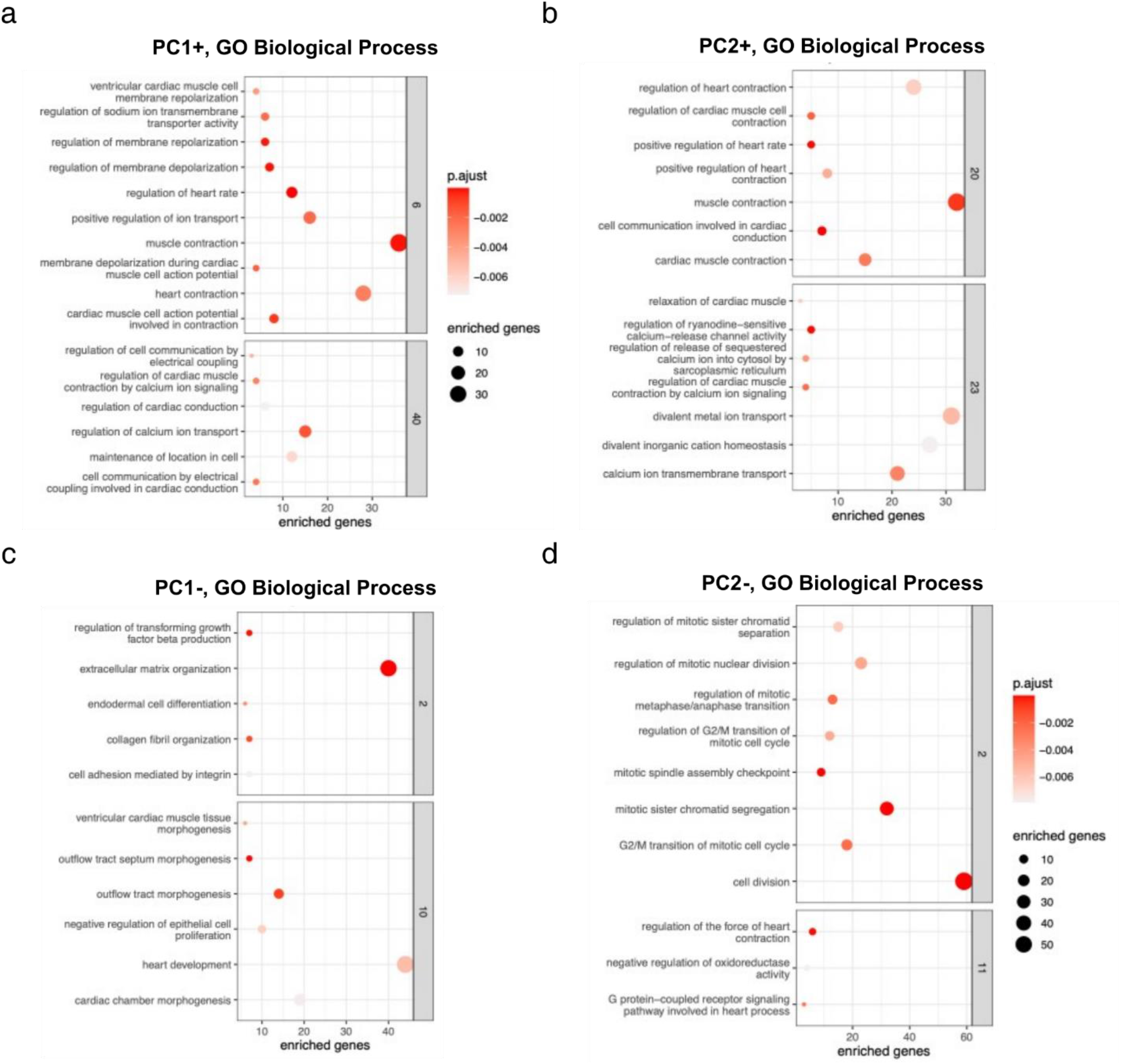
Meta-analysis of iPSC-CM maturation methods and *in vitro* myocardial development. Select clusters of enriched GO biological process terms corresponding to the indicated loadings within the principal component meta-analysis (**Figure 4l**) for PC1 **(a, c)** and PC2 **(b, d)**, positive **(a, b)**, and negative **(c, d)**. Point size denotes the size of the enriched gene set corresponding to each term. **Table S2** includes detailed descriptions and references for each category.

**Figure S6.**
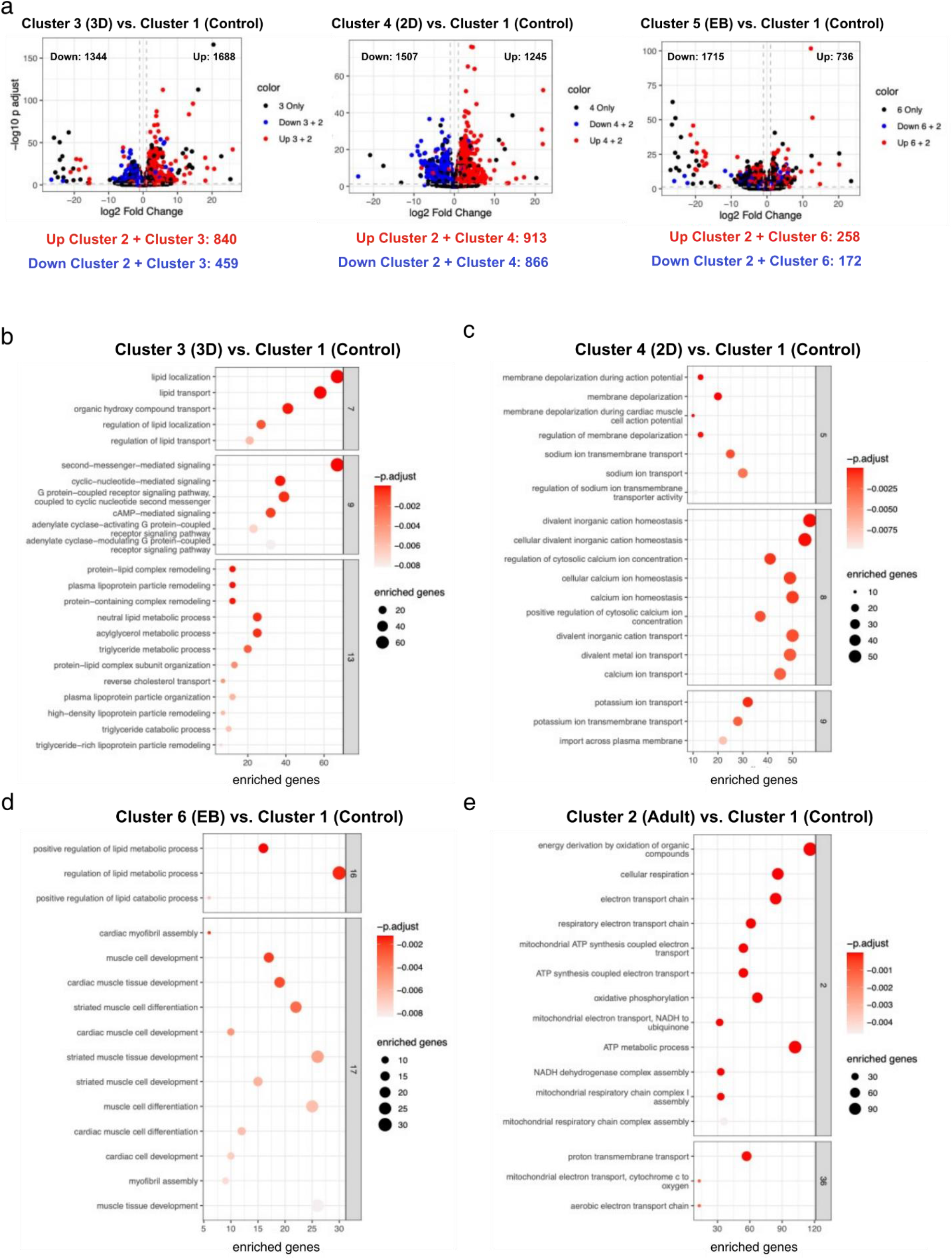
Cluster comparisons from meta-analysis of RNAseq datasets from PSC-CM cultures and in vivo patient samples. **a)** Volcano plots displaying log2 fold changes and log10 adjusted p-values for each gene in the indicated comparisons. Grey dashed lines denote the threshold for statistical significance (log2 fold change > ± 1 and p_adj_ < 0.05). Red and blue points represent genes significantly upregulated and downregulated, respectively, in adult vs. control. The number of significantly upregulated genes in the indicated comparison, and both the indicated comparison and adult vs. control are listed below the corresponding graph. **b-e)** Select clusters of enriched GO biological process terms in the indicated comparisons. Point size denotes the size of the enriched gene set corresponding to each term.

**Figure S7.**
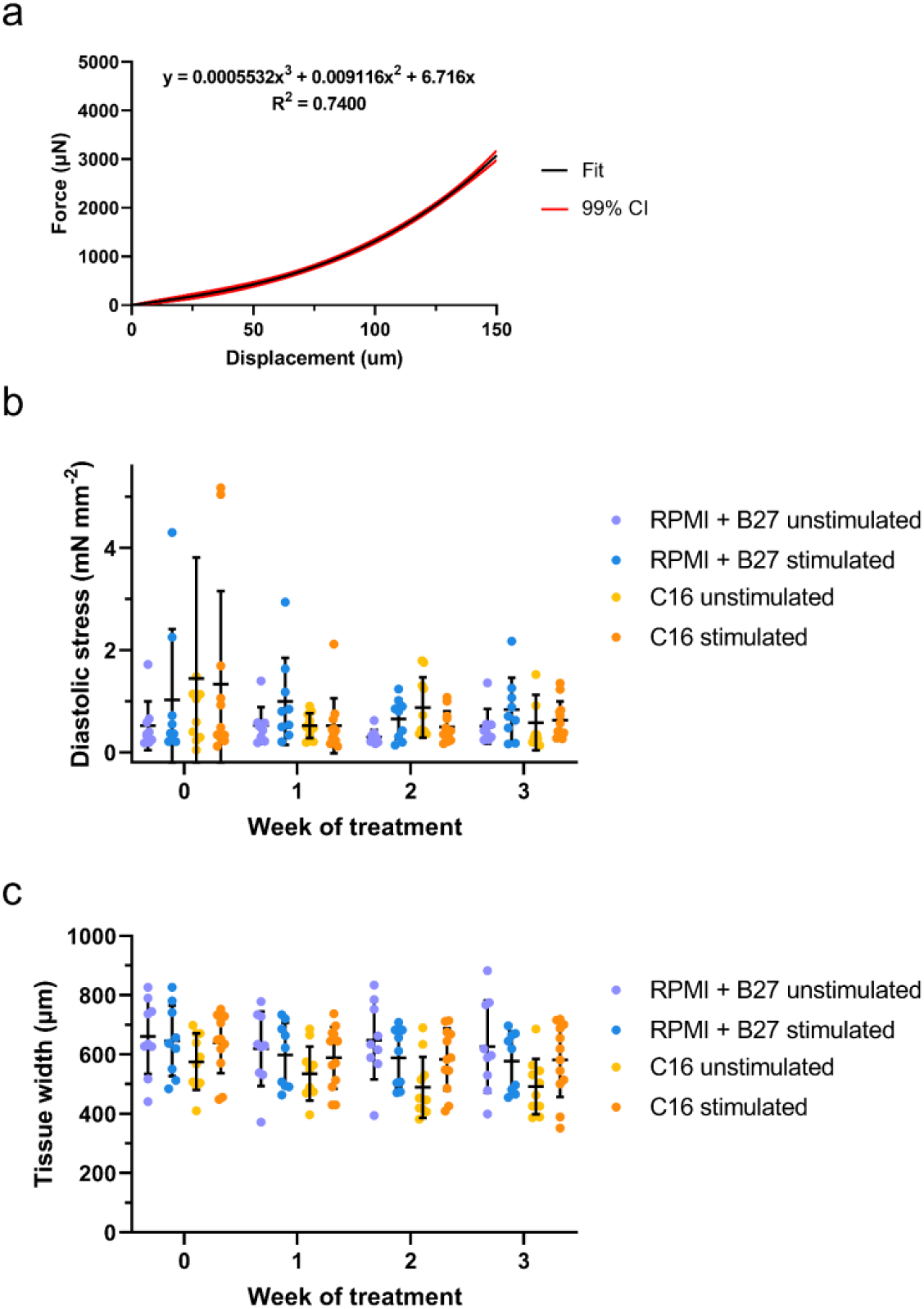
Calibration and characterization of maturing engineered myocardial tissues (Biowire II cultures). **a)** Compliance curves (cubic fit; 99% CI denoted by red borders) demonstrate the force-displacement relationship used to calibrate tissue deflection of polymeric beams to absolute forces. **b)** Diastolic stress (passive tension normalized to cross-sectional area) of tissues does not differ between medium treatments or stimulation conditions. **c)** Time dependence of tissue width (top-down view) of tissues.

**Figure S8.**
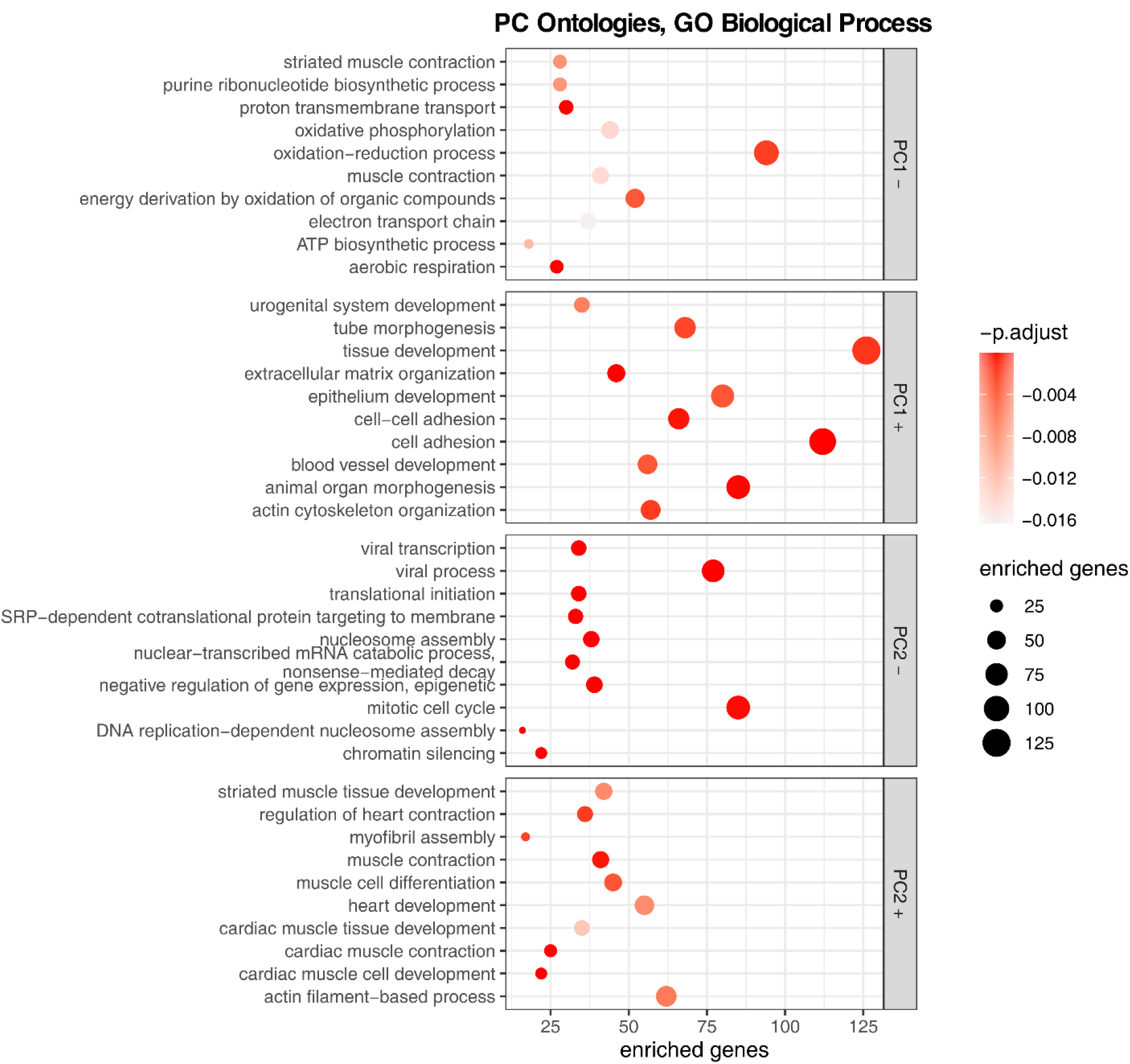
Transcriptionally-enriched functional terms and pathways in C16-treated Biowire II cultures. Top 10 enriched GO Biological Process terms for each direction and principal component in bulk RNA sequencing of Biowire II cultures treated with C16 or RPMI + B27 media, with or without electrical stimulation (**Figure 6i**). Point size denotes the number of significantly upregulated genes in each enriched category.

**Supplemental videos 1-4:** Representative Biowire II co-cultures contracting after 3 weeks in either C16 iPSC-CM maturation medium or RPMI + B27, with or without continuous electrical field pacing at 1 Hz. Tissues were spontaneously contracting for sample videos.

**Table S1.**
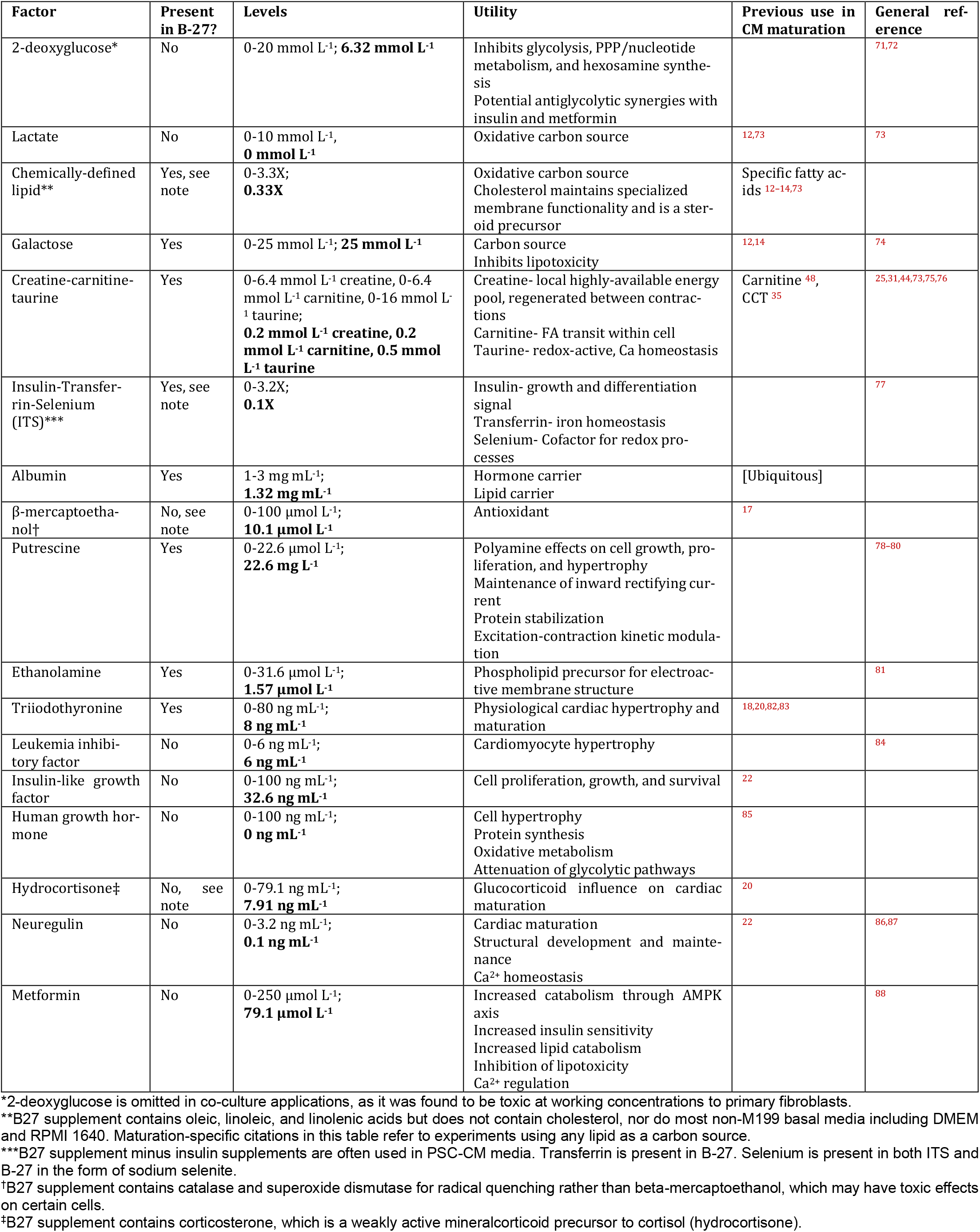
Soluble factors used in iteration for inclusion in an M199-based HD-DE screen, with individual putative physiological effects, as well as references to previous use specifically in PSC-CM maturation, or general physiological effects. The bolded level is the effective concentration in C16 medium; order corresponds to labels of **Figure 1c**. Numerous studies have used RPMI + B27 supplement ^13,15,17,23–26,35^; B27 additionally contains progesterone and pipecolic acid, which have no analogue in the compounds used in this study. Lactate and human growth hormone were not used in the final C16 formulation.

| Factor | Present in B-27? | Levels | Utility | Previous use in CM maturation | General reference |
| --- | --- | --- | --- | --- | --- |
| 2-deoxyglucose* | No | 0-20 mmol L<sup>-1</sup>; **6.32 mmol L<sup>-1</sup>** | Inhibits glycolysis, PPP/nucleotide metabolism, and hexosamine synthesis<br>Potential antiglycolytic synergies with insulin and metformin |  | <sup>71,72</sup> |
| Lactate | No | 0-10 mmol L<sup>-1</sup>, **0 mmol L<sup>-1</sup>** | Oxidative carbon source | <sup>12,73</sup> | <sup>73</sup> |
| Chemically-defined lipid** | Yes, see note | 0-3.3X; **0.33X** | Oxidative carbon source<br>Cholesterol maintains specialized membrane functionality and is a steroid precursor | Specific fatty acids <sup>12–14,73</sup> |  |
| Galactose | Yes | 0-25 mmol L<sup>-1</sup>; **25 mmol L<sup>-1</sup>** | Carbon source<br>Inhibits lipotoxicity | <sup>12,14</sup> | <sup>74</sup> |
| Creatine-carnitine- taurine | Yes | 0-6.4 mmol L<sup>-1</sup> creatine, 0-6.4 mmol L<sup>-1</sup> carnitine, 0-16 mmol L<sup>-1</sup> taurine; **0.2 mmol L<sup>-1</sup> creatine, 0.2 mmol L<sup>-1</sup> carnitine, 0.5 mmol L<sup>-1</sup> taurine** | Creatine- local highly-available energy pool, regenerated between contractions<br>Carnitine- FA transit within cell<br>Taurine- redox-active, Ca homeostasis | Carnitine <sup>48</sup>, CCT <sup>35</sup> | <sup>25,31,44,73,75,76</sup> |
| Insulin-Transfer- rin-Selenium (ITS)*** | Yes, see note | 0-3.2X; **0.1X** | Insulin- growth and differentiation signal<br>Transferrin- iron homeostasis<br>Selenium- Cofactor for redox processes |  | <sup>77</sup> |
| Albumin | Yes | 1-3 mg mL<sup>-1</sup>; **1.32 mg mL<sup>-1</sup>** | Hormone carrier<br>Lipid carrier | [Ubiquitous] |  |
| β-mercaptoethanol† | No, see note | 0-100 μmol L<sup>-1</sup>; **10.1 μmol L<sup>-1</sup>** | Antioxidant | <sup>17</sup> |  |
| Putrescine | Yes | 0-22.6 μmol L<sup>-1</sup>; **22.6 mg L<sup>-1</sup>** | Polyamine effects on cell growth, proliferation, and hypertrophy<br>Maintenance of inward rectifying current<br>Protein stabilization<br>Excitation-contraction kinetic modulation |  | <sup>78–80</sup> |
| Ethanolamine | Yes | 0-31.6 μmol L<sup>-1</sup>; **1.57 μmol L<sup>-1</sup>** | Phospholipid precursor for electroactive membrane structure |  | <sup>81</sup> |
| Triiodothyronine | Yes | 0-80 ng mL<sup>-1</sup>; **8 ng mL<sup>-1</sup>** | Physiological cardiac hypertrophy and maturation | <sup>18,20,82,83</sup> |  |
| Leukemia inhibitory factor | No | 0-6 ng mL<sup>-1</sup>; **6 ng mL<sup>-1</sup>** | Cardiomyocyte hypertrophy |  | <sup>84</sup> |
| Insulin-like growth factor | No | 0-100 ng mL<sup>-1</sup>; **32.6 ng mL<sup>-1</sup>** | Cell proliferation, growth, and survival | <sup>22</sup> |  |
| Human growth hormone | No | 0-100 ng mL<sup>-1</sup>; **0 ng mL<sup>-1</sup>** | Cell hypertrophy<br>Protein synthesis<br>Oxidative metabolism<br>Attenuation of glycolytic pathways | <sup>85</sup> |  |
| Hydrocortisone‡ | No, see note | 0-79.1 ng mL<sup>-1</sup>; **7.91 ng mL<sup>-1</sup>** | Glucocorticoid influence on cardiac maturation | <sup>20</sup> |  |
| Neuregulin | No | 0-3.2 ng mL<sup>-1</sup>; **0.1 ng mL<sup>-1</sup>** | Cardiac maturation<br>Structural development and maintenance<br>Ca<sup>2+</sup> homeostasis | <sup>22</sup> | <sup>86,87</sup> |
| Metformin | No | 0-250 μmol L<sup>-1</sup>; **79.1 μmol L<sup>-1</sup>** | Increased catabolism through AMPK axis<br>Increased insulin sensitivity<br>Increased lipid catabolism<br>Inhibition of lipotoxicity<br>Ca<sup>2+</sup> regulation |  | <sup>88</sup> |

\*2-deoxyglucose is omitted in co-culture applications, as it was found to be toxic at working concentrations to primary fibroblasts.
\*\*B27 supplement contains oleic, linoleic, and linolenic acids but does not contain cholesterol, nor do most non-M199 basal media including DMEM and RPMI 1640. Maturation-specific citations in this table refer to experiments using any lipid as a carbon source.
\*\*\*B27 supplement minus insulin supplements are often used in PSC-CM media. Transferrin is present in B-27. Selenium is present in both ITS and B-27 in the form of sodium selenite.
†B27 supplement contains catalase and superoxide dismutase for radical quenching rather than beta-mercaptoethanol, which may have toxic effects on certain cells.
‡B27 supplement contains corticosterone, which is a weakly active mineralcorticoid precursor to cortisol (hydrocortisone).

**Table S2.** Description of RNA-seq meta-analysis samples. “Adult”, “Infant”, and “Fetal” samples represent patient-derived myocardium or CM extracts thereof. “EB” samples represent PSC-CM differentiations and maturation in embryoid body format. “2D” samples represent matured PSCCM monolayers, while “3D” samples represent engineered myocardial tissues. “Control” samples represent differentiated but unmatured PSC-CMs (*i*.*e*., harvested at the point of induction of maturation treatments).

| Batch | Type | Description | Reference | GEO Accession |
| --- | --- | --- | --- | --- |
| A | Control | Day 20 immature PSC-CM | 52 | GSE62913 |
|  | Fetal | Fetal cardiomyocyte, *in vivo* |  |  |
|  | Adult | Adult cardiomyocyte, *in vivo* |  |  |
|  | 2D | PSC-CM in monolayer + let-7 overexpression |  |  |
| B | 3D | PSC-CM in ventricular Biowire, stimulated | 43 | GSE114976 |
|  | 3D | PSC-CM in ventricular Biowire, non-stimulated |  |  |
| C | 2D | PSC-CM in monolayer + fatty acids | 48 | GSE124057 |
| D | Control | Day 20 immature PSC-CM | 35 | GSE151279 |
|  | 2D | PSC-CM in monolayer + complex defined maturation medium |  |  |
| E | 2D | PSC-CM in monolayer + 90 day culture period | 53 | GSE81585 |
|  | Adult | Adult cardiomyocyte, *in vivo* |  |  |
| F | 3D | PSC-CM in Heart-Dyno | 13 | GSE62913 |
|  | Control | Adult cardiomyocyte, *in vivo* |  |  |
| G | 3D | PSC-CM in organoid co-culture with fibroblasts/endothelial cells | 49 | GSE116464 |
|  | Control | Day 20 immature PSC-CM |  |  |
| H | Fetal | Fetal cardiomyocyte, *in vivo* | 54 | PRJNA353755 |
|  | Adult | Adult cardiomyocyte, *in vivo* |  |  |
|  | Infant | Infant cardiomyocyte, *in vivo* |  |  |
| I | EB | PSC-CM in EB + glucose deprivation | 50 | GSE84815 |
|  | Control | Day 20 immature PSC-CM |  |  |
| J | EB | PSC-CM in EB + complex defined maturation medium | 51 | GSE125862 |

